# Topography of putative bidirectional interaction between hippocampal sharp wave ripples and neocortical slow oscillations

**DOI:** 10.1101/2024.10.23.619879

**Authors:** Rachel Swanson, Elisa Chinigò, Daniel Levenstein, Mihály Vöröslakos, Navid Mousavi, Xiao-Jing Wang, Jayeeta Basu, György Buzsáki

## Abstract

Systems consolidation relies on coordination between hippocampal sharp-wave ripples (SWRs) and neocortical UP/DOWN states during sleep. However, whether this coupling exists across neocortex and the mechanisms enabling it remain unknown. By combining electrophysiology in mouse hippocampus (HPC) and retrosplenial cortex (RSC) with widefield imaging of dorsal neocortex, we found spatially and temporally precise bidirectional hippocampo-neocortical interaction. HPC multi-unit activity and SWR probability was correlated with UP/DOWN states in mouse default mode network, with highest modulation by RSC in deep sleep. Further, some SWRs were preceded by the high rebound excitation accompanying DMN DOWN→UP transitions, while large-amplitude SWRs were often followed by DOWN states originating in RSC. We explain these electrophysiological results with a model in which HPC and RSC are weakly coupled excitable systems capable of bi-directional perturbation and suggest RSC may act as a gateway through which SWRs can perturb downstream cortical regions via cortico-cortical propagation of DOWN states.

---

Theories of systems consolidation rely on hippocampal-mediated coordination of neural activity across neocortex in service of reactivation during sleep [1]–[5]. However, how and to what extent this spontaneously occurs across regions, often many synapses removed from the hippocampus, remains unknown. During NREM sleep, neural populations alternate between periods of spiking and inactivity, termed UP and DOWN states in the neocortex, and sharp wave-ripples (SWRs) and inter-SWRs (iSWRs) in the hippocampus. Both gain and loss of function studies demonstrate the importance of the tight temporal coordination of these events for systems consolidation [6], [7]. However, the observed timing of this coordination is variable across experiments and regions, leading to a lack of mechanistic consensus regarding the inter-regional interaction required for consolidation.

Most studies agree that the probability of SWRs is higher during UP states and that the spike content of SWRs is biased by neocortical inputs [8]–[13], but see [14]–[16]. Some studies further suggest that SWRs initiate neocortical UP states [14], [17], [18], while others, in contrast, indicate that DOWN states follow SWRs [11], [12], [19]. These discrepancies may be due to variation in sleep depth, which modulates the rate of both SWRs and DOWN states [14], [20]–[23], or differences between cortical regions, especially given that UP/DOWN states can be localized [24] or travel across the forebrain [25], [26].

In an attempt to resolve these ambiguities, imaging studies have explored the topographic relationship between SWRs and the rest of the brain. In primates, SWRs were correlated with an increase in the BOLD signal in regions comprising the default-mode network (DMN; [23], [27]), similarly observed in humans using MEG [28]. Although of functional interest given the importance of the DMN for episodic recall [29], [30], only recently have rodent widefield imaging studies had the spatiotemporal resolution necessary to explore short timescale interaction between the hippocampus and dorsal neocortex, but with variable results [31]–[33]. Thus, where, when, and how SWRs are coupled with neocortical UP/DOWN states remains an unresolved tension across theories of systems consolidation.

Towards this goal, we developed a chronic preparation in mice that combined widefield imaging in the dorsal neocortex with silicon probe recordings of hippocampus and RSC in the same hemisphere. We found a topographically specific, state-dependent, bi-directional interaction between hippocampal SWRs and neocortical UP/DOWN states. From the neocortex to the hippocampus, SWRs were less likely to occur during DOWN states across regions in the default mode network, and SWRs often followed large rebound excitation at the DOWN-UP transition in DMN. From the hippocampus to the neocortex, large amplitude SWRs were often followed by DOWN states in RSC and motor cortical regions that then propagated along dorsal neocortex. The highest modulation was seen in RSC during deep NREM sleep in all cases. We hypothesized that these experimental observations could arise from weakly coupled populations in the complementary excitable regimes characteristic of NREM [34], and confirmed the plausibility of this hypothesis with a mean-field model of bi-directionally interacting hippocampal and RSC populations.

## RESULTS

### Combined wide-field imaging and chronic extracellular electrophysiology for studying hippocampal-cortical interaction during sleep

We combined chronic electrophysiological recordings from the hippocampus (HPC) and retrosplenial cortex (RSC) with widefield imaging of the dorsal neocortex in head-fixed Thy1 GCaMP6f mice (**Fig. 1A**; [35]). To record concurrently in the same hemisphere, a single-shank silicon probe (64 or 128 recording sites) was lowered through the left hemisphere to the right RSC and hippocampal CA1 regions, ipsilateral to our thinned-skull cranial window preparation (**Fig. 1A-E**). Following hemodynamic correction ([36]; see **Methods**) and alignment of widefield videos to the Allen Institute’s Common Coordinates Framework (**Fig. S1**; [37]; see **Methods**), we confirmed the successful placement of our recording electrode by verifying that the correlation between extracellularly recorded RSC population rate and all widefield pixels was highest in RSC (**Fig. 1B**, red dots). To recover fine timescale changes in population rate across our imaging field of view, we determined a deconvolution kernel that optimally predicted electrically recorded RSC population rate from the identified RSC region of interest (ROI) in each mouse (**Fig. S2; Suppl. Movie 1**; [38]; see **Methods**). We next deconvolved widefield activity across neocortex for each mouse with the derived kernel, as was successfully done previously [38]. Variation in standard deviation of deconvolved pixel time series across regions was minimal (**Fig. S2**). The remaining analyses were performed with either deconvolved widefield activity or unaltered fluctuations in total blood volume (Hemoglobin Hbt; 525 nm), as specified. This approach uniquely combined optical measurement of the population rate of excitatory cells across the dorsal neocortical mantle (**Fig. 1B-C**) with simultaneous extracellular recordings in the hippocampus and RSC in the same hemisphere (**Fig. 1D-E**).

**Figure 1.**
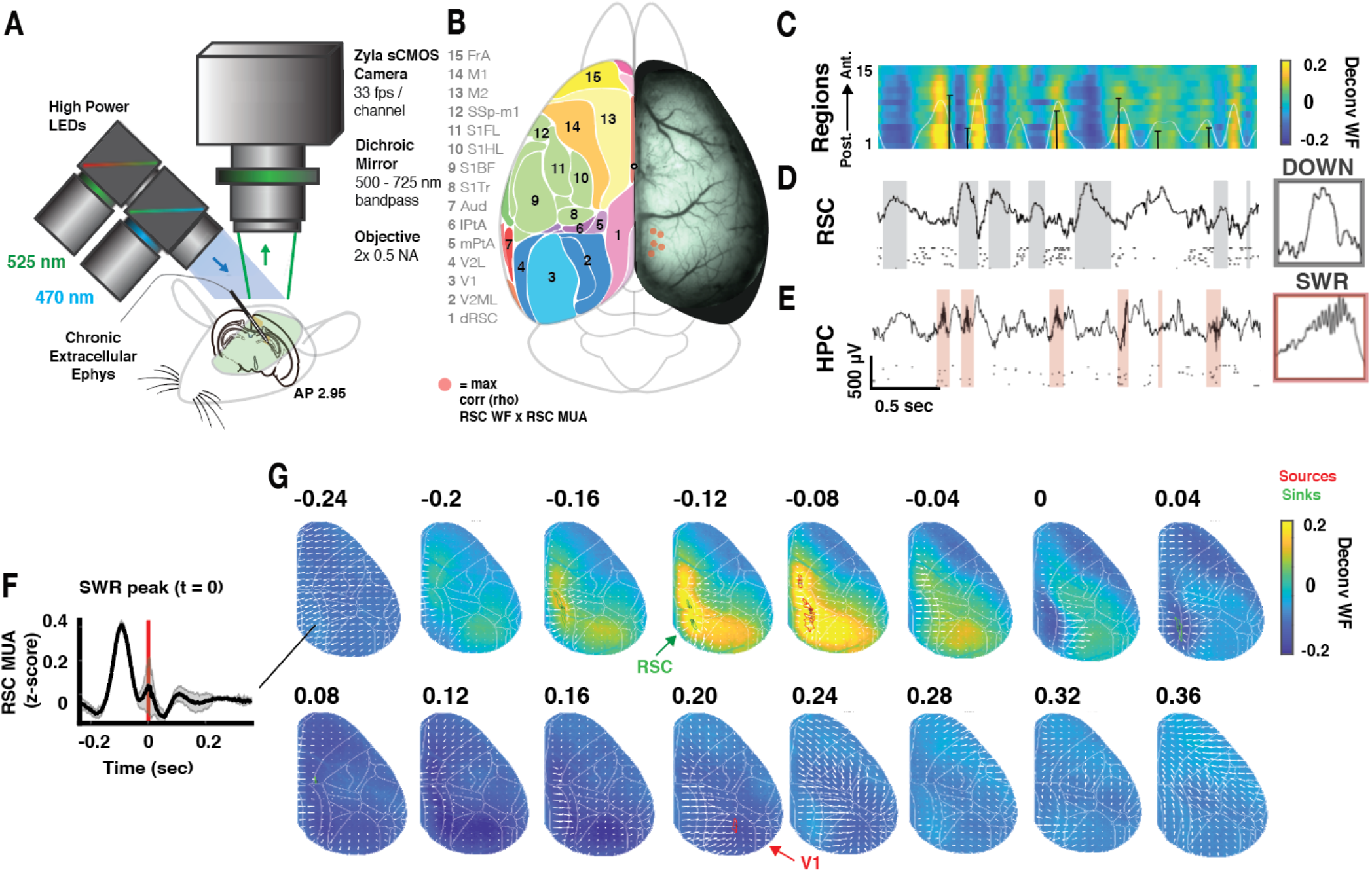
Experimental preparation and neocortical activity surrounding hippocampal SWRs. **A**. Dual wavelength (blue 470 nm – thy1 GCaMP6f; green 525 nm – total blood volume) widefield imaging (66 frames per second) of the dorsal hemisphere of a thy1 GCaMP6f mouse. Note chronic silicon probe spanning ipsilateral CA1 and RSC beneath the imaging field of view (green). **B**. *Right*, Example raw fluorescence frame. *Left*, Corresponding cortical regions. Red dots indicate location of maximum correlation (rho) between widefield signal and RSC population rate for each mouse (n=5). **C-E**. Aligned simultaneous widefield imaging of dorsal cortex and electrophysiological recordings in HPC and RSC. **C**. Deconvolved widefield time series for 15 pixels in regions ranging from posterior to anterior dorsal cortex as in B. White line corresponds to RSC widefield time series (also row 1 in heat map); black bars denote SWRs, height proportional to SWR amplitude. **D-E**. Example LFP and single units from RSC and hippocampal CA1 pyramidal layer. Shaded areas highlight DOWN states and SWRs in RSC and HPC, respectively. Right insets, example DOWN state and SWR (100 ms). **F**. Average RSC multi-unit activity (MUA; see **Methods**) surrounding all SWR peaks at t = 0. Shading corresponds to standard deviation across mice (n = 5). **G**. Average deconvolved widefield activity across all mice surrounding SWR peak at t = 0. Sources and sinks are identified in green and red, respectively. Arrows correspond to vector fields calculated across pairs of frames on the grand-average video, providing a qualitative view of activity flow.

As observed electrophysiologically (**Fig. 1F**; peak time t = 0, cites), SWRs were preceded by elevated neocortical activity in the deconvolved widefield data (**Fig. 1G, Suppl. Movie 2**), led by a source in RSC (t = –0.12 s) that spread throughout midline-posterior cortical regions (mouse DMN or “medial networks” [39]). This increased activity was followed by decreased activity in RSC that spread across the neocortex, ultimately terminating with a sink in V1 (t = 0.2 s).

### Joint fluctuation of SWRs and cortical DOWN states across ultraslow (0.01 – 0.03 Hz), infraslow (0.04 – 0.5 Hz), and slow (0.5 – 4 Hz) timescales

Next, we examined whether hippocampal-cortical coupling varied as animals shifted from wake to sleep. Automated classification of brain states was performed using three variables: the time-varying slope of the RSC power spectrum (power spectral slope, PSS); [40]), HPC theta power, and LFP-derived electromyogram (pseudo-EMG) (**Fig. 2A-B**; [20], [41]). This resulted in 3 clusters that corresponded to active WAKE (high EMG), REM, and a third cluster that ranged from quiet WAKE (low EMG) to NREM (**Fig. 2B**). To ensure that the brain states observed during head-fixation were comparable to natural behavior, we state-scored concatenated head-fixed and home cage recording sessions within the same mouse (**Fig. 2A-B**). While the fraction of time spent in each state varied between conditions, the regular recurrence of transitions from deep NREM to REM sleep in both conditions and the qualitatively overlapping head-fixed and home cage brain state clusters confirmed comparable sleep quality in head-fixed animals (**Fig. S3** for individual mice).

**Figure 2.**
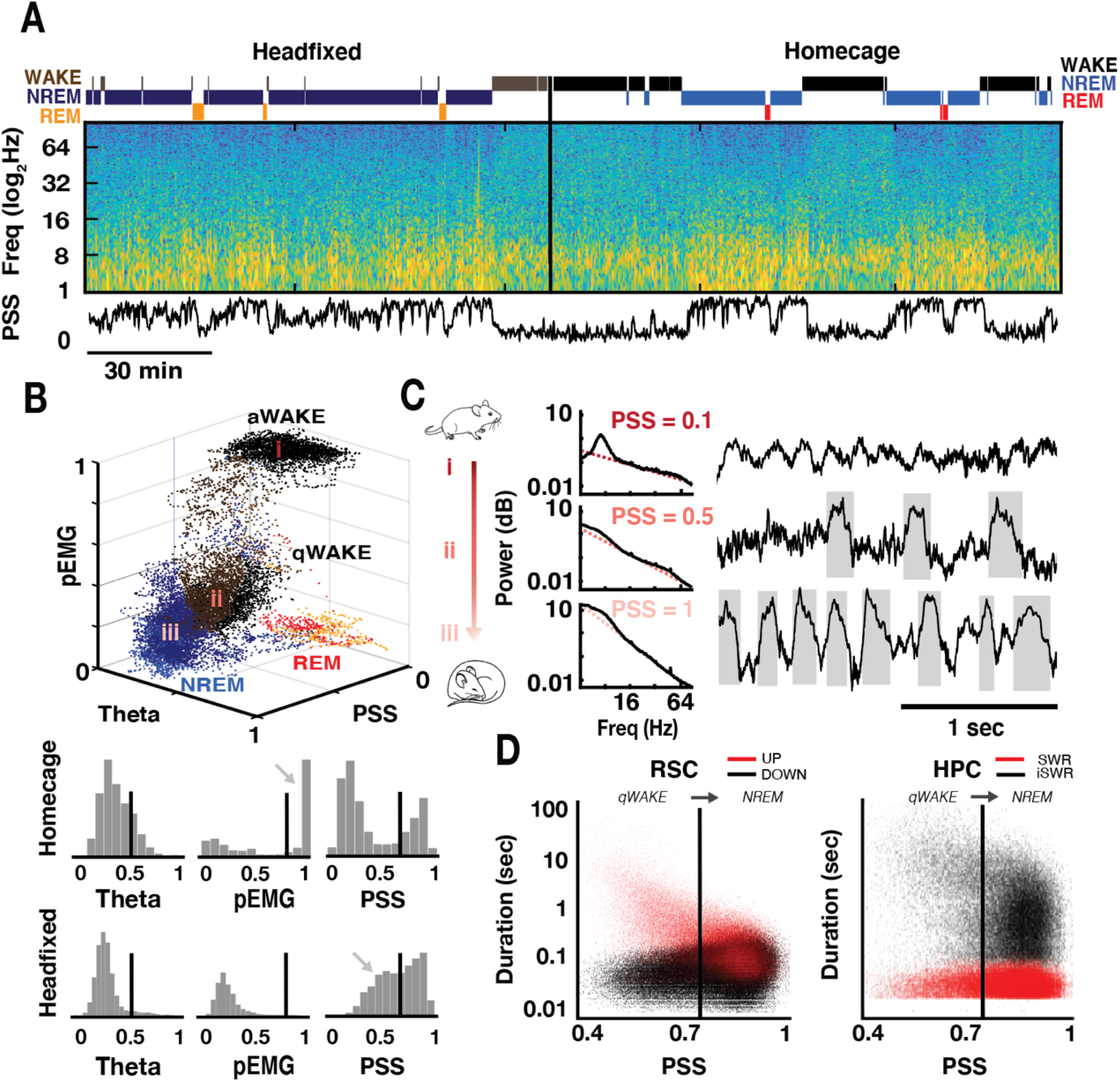
SWR and DOWN state rates increase as animals move from quiet wake to deep NREM sleep. **A**. Brain state-scoring of concatenated headfixed and home cage recording sessions for an example mouse. *Top*, Identified WAKE, NREM, and REM states. *Middle*, spectrogram of RSC LFP. *Bottom*, Time-varying slope of the power spectrum (PSS). **B**. *Top*, State scoring of the session in panel A. Note three distinct clusters, classified as active wake (aWAKE), REM sleep, and a third cluster with continuous variation from quiet wake (qWAKE) to NREM sleep. *Bottom*, Distributions of the three variables used for behavioral state scoring (PSS, proxy EMG, and theta power) in homecage and headfixed conditions. **C**. Average RSC power spectra (black; left) and example RSC LFP traces (right) at three different arousal levels from active WAKE to deep NREM, denoted i-iii in panel B scatterplot. Inset PSS values are the inverse of the slope of the linear fit to the aperiodic component of the power spectra (pink dotted lines). DOWN states are shaded in gray. **D**. *Left*, Scatter plot of durations of UP (red) and DOWN (black) states in RSC across values of PSS for all mice. *Right*, Scatter plot of dwell time durations for SWRs (red) and inter-SWR periods (black). Vertical lines in RSC and HPC separate qWAKE and NREM.

Hippocampal SWRs and RSC UP/DOWN states were observed exclusively throughout the brain state cluster comprised of quiet WAKE and NREM sleep (labeled ii and iii in **Fig. 2B**). However, their frequency of occurrence varied continuously as a function of PSS, or arousal level (**Fig. 2C**; [42]). From quiet WAKE (low PSS) to deep NREM sleep (high PSS), DOWN state rate increased (**Fig. 2C**). This occurred because the duration of RSC UP states got increasingly shorter (**Fig. 2D**, left red) and the duration of DOWN states became increasingly more variable (**Fig. 2D**, left black), until the ratio of mean UP and DOWN state durations approached one. The hippocampus followed a complementary pattern: as PSS values increased, the rate of hippocampal SWRs increased due to a decrease in the inter-SWR interval (**Fig. 2D** right black).

Hippocampal SWRs were further modulated by RSC UP and DOWN states, with SWRs significantly more likely during UP states (**Fig. 3A**). This relative change in SWR rate from DOWN to UP states increased monotonically with increasing PSS, ultimately resulting in a 3-fold increase in SWR rate from DOWN to UP states during deep NREM (**Fig. 3B-C**), parallel with increased RSC multi-unit activity, MUA, within UP states (**Fig. 3D**). Hippocampal MUA likewise increased with increasing RSC UP state firing rate, following RSC D-U transitions with a time lag despite a near-synchronous decrease in RSC rate at the U-D transition (**Fig. 3E**; ‘co-active and co-silent frames’; [9], [10], [15]). In sum, the modulation of hippocampal activity by RSC UP/DOWN states depended on arousal level, as measured by PSS. With decreasing arousal, the mean firing rate of RSC UP states increased and was paralleled by an increase in HPC MUA and subsequent increased probability of SWRs.

**Figure 3.**
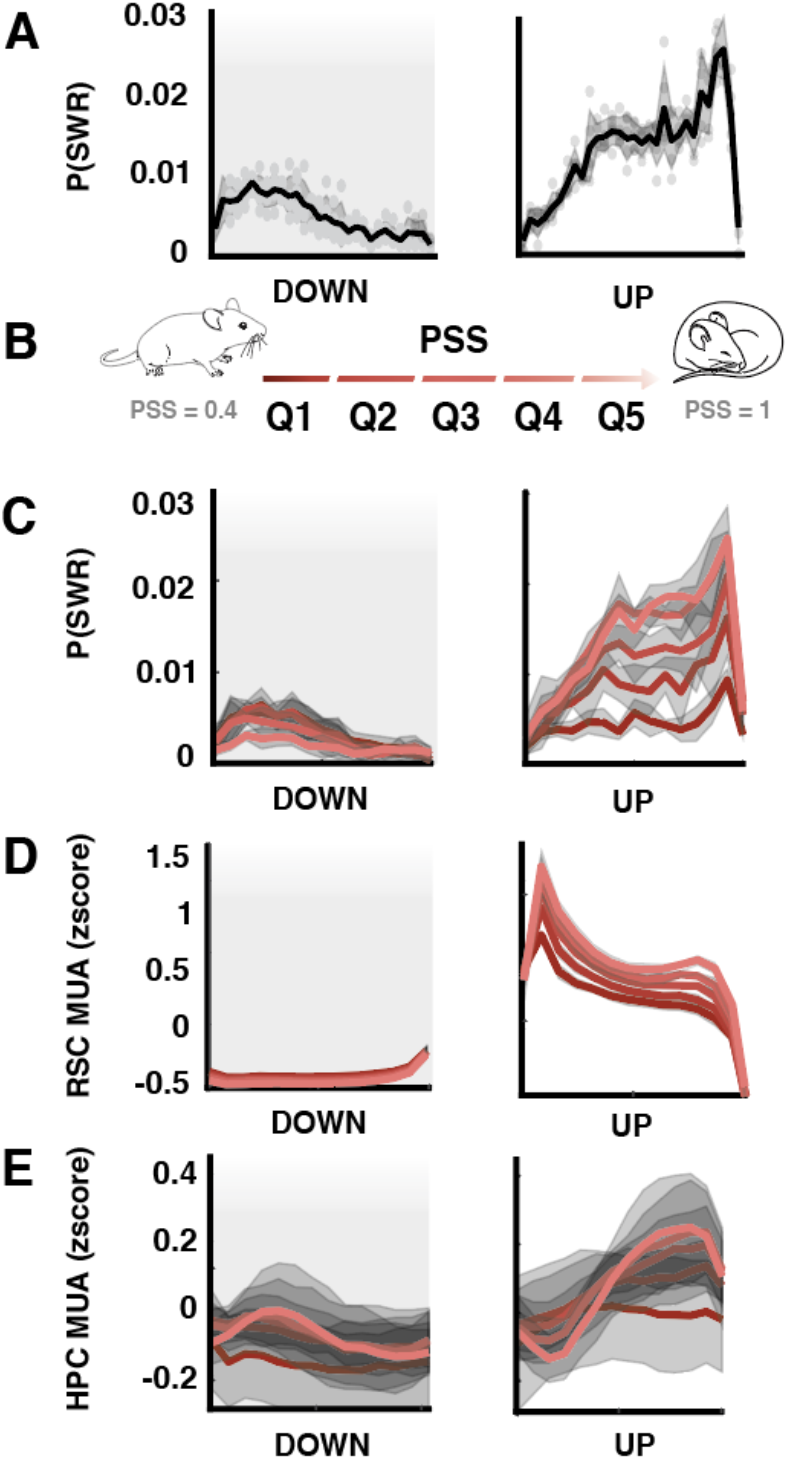
RSC UP and DOWN states modulate hippocampal SWRs as a function of brain state. **A**. Probability of SWRs across time-normalized RSC UP and DOWN states. Shading corresponds to standard deviation across mice; dots to individual mice. **B**. PSS quintiles span quiet WAKE to deep NREM (Q1-Q5; colored from dark to light red in all panels). **C-E**. Variables specified plotted across time-normalized RSC UP and DOWN states as a function of PSS quintile; all mice. Shading corresponds to standard deviation across all UP or all DOWN states. **C**. Probability SWR by PSS quintile. **D**. Mean RSC MUA by PSS quintile. **E**. Mean HPC MUA by PSS quintile.

Brain state, as measured using a variety of metrics, is known to fluctuate in both the “ultraslow” (0.01-0.03 Hz) and “infraslow” (0.04-0.5 Hz) frequency bands ([43]–[47], apparent in cortical blood flow [48], [49]. Enabled by green wavelength (525 nm) imaging of total blood volume (Hbt) across the neocortical mantle, we found that fluctuations in Hbt showed a 1/f background with peaks in “ultraslow” and “infraslow” frequency ranges (**Fig. S4**; **Suppl. Movie 3**). Variation in PSS more closely tracked fluctuation in the ultraslow-filtered Hbt (**Fig. S4D**), which was globally coherent across the cortical mantle (**Fig S4E**). In contrast, the infraslow-filtered Hbt was accompanied by a faster-timescale modulation of SWR rate, confined to the DMN (**Fig. S4F**; [28]). This phase-dependence was not restricted to SWRs, but rather reflected a broader infraslow-timescale switch in RSC and HPC LFP between power spectra typical of NREM to a state dominated by 4 Hz in RSC (**Fig. S4G**).

Together, these results reveal co-modulation of hippocampal-cortical state at three timescales: 1) an ultraslow (0.01-0.03Hz) variation in brain state (perhaps analogous to the ‘global signal’ in fMRI ([50], [51]), measured by the time-varying slope of the power spectrum (PSS) and fluctuations in total blood volume (Hbt), and accompanied by concurrent changes in the rate of DOWN states, SWRs, and cortical spiking activity during UP states, 2) an infraslow (0.04-0.5Hz) fluctuation of cortical state in mouse default mode network (perhaps reflecting excitability changes during NREM sub-stages, or “packets” [20], and 3) a slow (.5 – 4 Hz) modulation of SWR rate by RSC UP and DOWN states.

### Putative bidirectional hippocampal-cortical perturbation by transient population synchrony

Motivated by previously observed temporal coupling between SWRs and cortical slow waves [12], [52], and the finding that SWRs cluster towards the end of time-normalized UP states (**Fig. 3A, C**), we next investigated whether UP/DOWN state transitions in RSC could predict the timing of SWRs. When aligned to DOWN to UP (D-U) or UP to DOWN (U-D) transitions (**Fig. 4B-D, Fig. S5**), RSC MUA was asymmetric, displaying a peak at the D-U transition not present at the U-D transition (putative K-complex, K; **Fig. 4D**). In parallel, we observed a tight clustering of SWRs around U-D and D-U transitions, with probability of SWR occurrence (pSWR) exhibiting three distinct peaks (**Fig 4D-E, S5**). First, a peak in pSWRs occurred within a 50ms time window prior to the U-D transition (SWRUD). Second, pSWR peaked within ∼80 ms after the U-D transition (SWRD). Finally, a peak in pSWR occurred after a ∼120ms delay after the D-U transition in RSC (SWRDU), following the D-U peak in RSC MUA. There were many more U-D and D-U state changes than the number of SWRs, so these hypothesized interactions took place during only a small fraction of cortical transitions. Nevertheless, more than half of the SWRs were time-locked to RSC D-U or U-D state transitions (SWRUD, SWRDU and SWRD types; **Fig. 4I**; **Fig. S6**). While SWR bursts (defined as inter-SWR interval of 50 - 132 ms) comprised only a small fraction (<20%) of all SWRs, burst onsets were more likely following the D-U transition (SWRDU), and burst offsets were more likely at the U-D transition (SWRUD), particularly surrounding long DOWN states (**Fig. S6**). These observations cannot simply be explained by tonic modulation of SWR rate by UP states, as UP state probability is symmetric surrounding U-D and D-U transitions (**Fig. 4B**).

**Fig 4.**
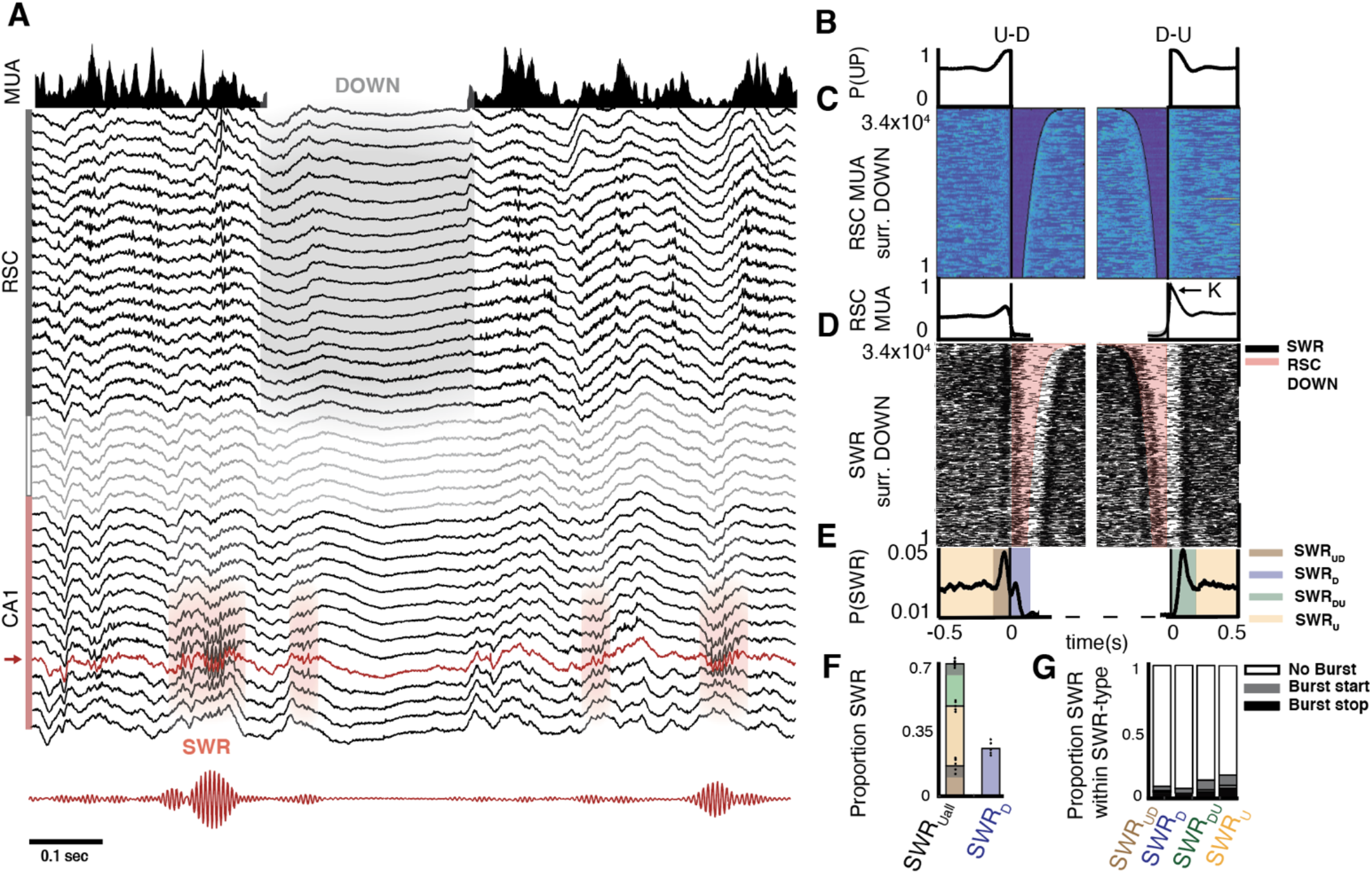
Probability of SWRs around UP-DOWN (U-D) and DOWN-UP (D-U) transitions is asymmetric. **A**. Example LFP traces spanning layers of granular RSC, white matter, and ipsilateral CA1; RSC MUA (above); ripple frequency filtered CA1 trace (below; 130-200 Hz; bandpass filtered channel designated in red). **B-E**. Data specified surrounding all DOWN states for an example mouse, centered at RSC U-D transitions (left) or D-U transitions (right) and sorted by DOWN state duration. **B**. Probability of being in an UP state, surrounding transitions. **C**. RSC MUA; each row is an U-D (left) or D-U (right) transition (>30,000). *Bottom*, average RSC MUA surrounding transition specified. K refers to transient rebound population synchrony at the D-U transition, K-complex or ‘K’. **D**. Raster plot of all SWRs during the same RSC U-D and D-U transitions as in **C**. Pink shading corresponds to RSC DOWN states identified in panel C. SWRs plotted as thin black lines, the length of which corresponds to their durations. Note decreased P(SWR) during DOWN, asymmetry in clustering of SWRs around transitions, and change in clustering as DOWN duration increases. **E**. Defining SWRs by their temporal proximity to U-D and D-U transitions yields 4 “types”, SWRU (yellow), SWR UD (red), SWR D (blue), and SWR DU (green); see **Methods** and **Fig. S6. F**. Proportion of each “SWR type” across all mice (dots represent individual mice; colors correspond to SWR type). Note 3-fold increase in SWR rate from DOWN to UP states. Gray shaded region in SWRUD and SWRDU represents the overlap between these categories (∼30%). **G**. For each SWR type, proportion of those SWRs that occur in bursts vs not in bursts (see Methods). Start and end times of the burst are denoted by gray and black.

The clustering of SWRs around U-D and D-U transitions suggests a more temporally precise, and potentially causal hippocampal-neocortical interaction; whereby hippocampal SWRs may induce U-D transitions in the cortex and the transient elevation of cortical MUA at D-U transitions (K-complex) may induce SWRs in the hippocampus (**Fig. 5A**; [15], [18], [34]). To test this possibility further, we examined the change in the probability of RSC DOWN states as a function of SWR amplitude (**Fig. 5B**), and the change in the probability of SWRs as a function of K-complex magnitude, defined as average RSC MUA within a 20ms window following the D-U transition (**Fig. 5E**). As the amplitude of SWRs increased, they were more likely to be followed by an U-D transition at a fixed 30 ±15 ms delay (**Fig. 5B**). The consistency of this lag suggests it is the time window in which hypothesized SWR-induced DOWN states occur. Similarly, as MUA at the D-U transition increased, the probability of SWRs increased at a fixed lag of 120±15 ms (**Fig. 5E**), suggesting the lag at which k-complex induction of SWRs may occur.

**Figure 5.**
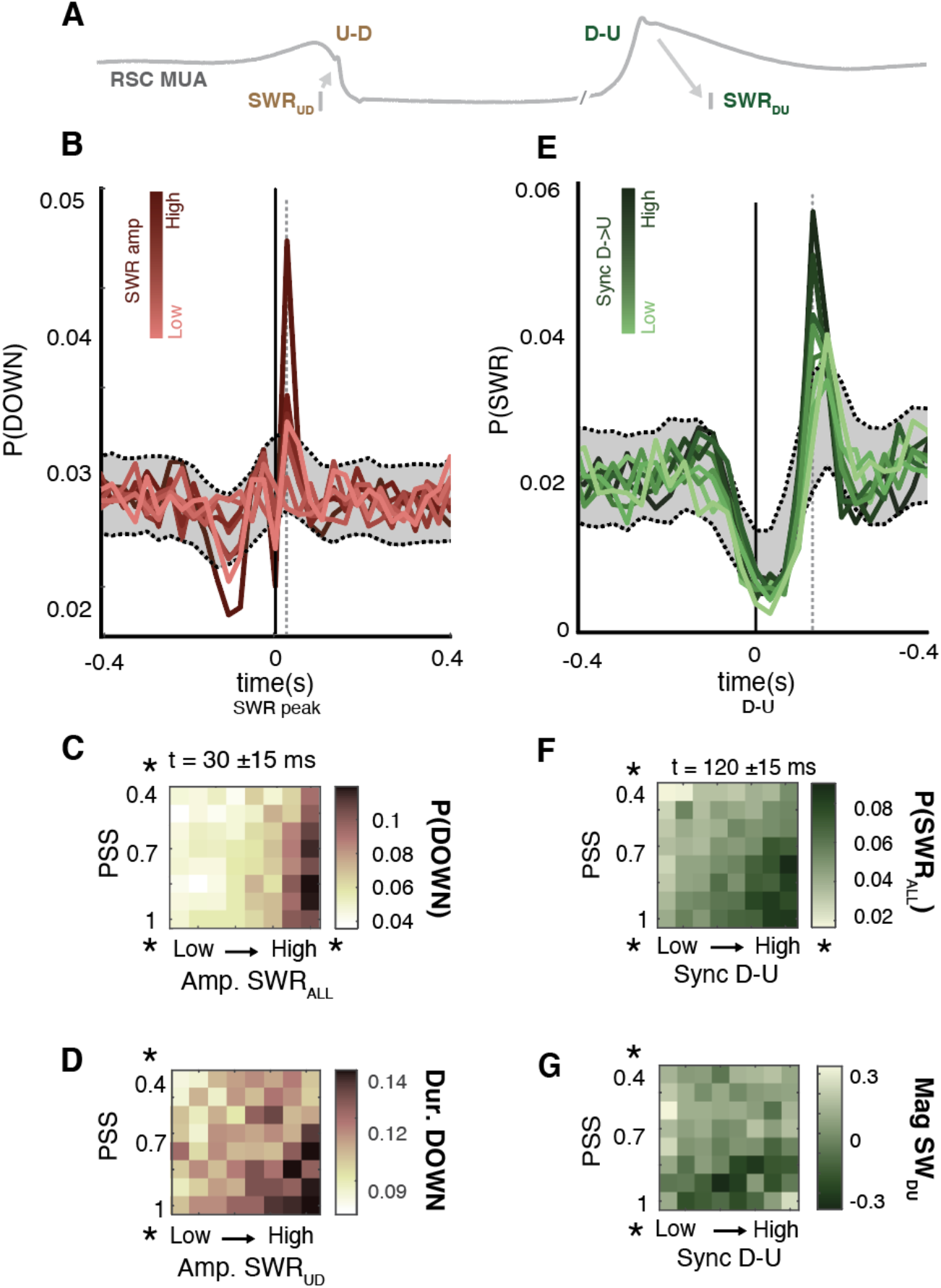
Temporal relationship between HPC and RSC state transitions is state-dependent and bi-directional. **A**. Schematic of hypothesis: SWRs can induce U-D transitions and D-U transitions can induce SWRs, conditional on magnitude of the perturbation and state of the receiving region. **B**. Cross-correlograms between SWR peaks (t = 0 s) and DOWN state onsets across all mice, colored by SWR amplitude octile (light to dark red; small to large SWRs). Shading denotes boot-strapped 99% confidence intervals obtained by shuffling both SWRALL peak and U-D time series by ±30ms, 1000 iterations. Note increased probability of DOWN onset at fixed 30 ± 15 ms timelag (vertical gray line) with increasing SWR amplitude. **C**. Mean probability of DOWN state onset at a 30ms lag from SWR peak, timelag of putative ‘interaction’, as a function of depth sleep (PSS) and SWRALL amplitude (repeated measures two-way ANOVA across sessions (n=15): R2 = 0.47. *SWR amplitude*, F = 83.19, p < 0.001, η^2^p = 0.42; *PSS*, F = 5.87, p < 0.001, η^2^p = 0.07; *Interaction*, F = 1.68, p < 0.05, η^2^p = 0.06). Significant effect of amplitude SWR, depth sleep, and their interaction. **D**. Mean duration of DOWN states following SWRUD as a function of depth sleep (PSS) and SWRUD amplitude across all mice (GLM 5-fold CV: R2 = 0.014. *SWR amplitude* β1 = –0.007, t = 0.006, p = *NS*; *PSS* β1 = 0.067, t = 7.68, p < 0.001; *Interaction* β1 = –0.016, t = 1.96, p < 0.05). **E**. Probability of SWRs surrounding RSC D-U transitions (t = 0s), colored by D-U rebound excitation octile (light to dark green, small to large). Note increase in P(SWR) with increasing rebound excitation at a fixed lag of 120ms (vertical gray line). Confidence intervals computed as in B. **F**. Mean probability of SWR occurrence at a 120ms lag from RSC D-U as a function of depth sleep (PSS) and D-U rebound excitation (repeated measures two-way ANOVA: R2 = 0.58. *Rebound excitation*, F = 54.01, p < 0.001, η^2^p = 0.32; *PSS*, F = 120.26, p < 0.001, η^2^p = 0.42; *Interaction*, F = 3.78, p < 0.001, η^2^p = 0.15. **K**. Mean magnitude of HPC sharp-waves as a function of tonic MUA HPC and D-U rebound excitation across all mice (GLM 5-fold CV: R2 = 0.05. *Rebound excitation* β1 = -0.27, t = -1.65 p = *NS*; *PSS* β1 = -1.01, t = -5.02, p < 0.001; *Interaction* β1 = 0.4, t = 1.95, p < 0.05).

We further found that the interaction between SWRs and UP-DOWN states was modulated by arousal level, as measured by PSS. Large amplitude SWRs were more likely to be followed by DOWN states in deep NREM (high PSS), with a significant effect of SWR amplitude, PSS, and their interaction (**Fig. 5C**). In addition, we found a significant effect of arousal level and the interaction of arousal level with SWR amplitude on DOWN duration (**Fig. 5D**), implying the duration of DOWN states is conditional on depth sleep and providing further support for a potential role of SWRs in DOWN state induction. Similarly, K-complex magnitude increased the probability of SWRs at a fixed lag of 120±15 ms, with a significant effect of magnitude K-complex, PSS, and their interaction (**Fig. 5F**). Further, the magnitude of sharp wave sink in stratum radiatum, a measure of the input drive to CA1 from CA3, became increasingly negative (corresponding to a larger sink) as a function of PSS and interaction of PSS with K-complex magnitude (**Fig. 5G**). Overall, these findings support the hypothesis that large amplitude SWRs may trigger U-D transitions (SWRUD) and that transient spike synchrony at D-U transitions (K-complex) may trigger SWRs (SWRDU and a fraction of SWRD when UP state is short; <100 ms). In both directions, the effectiveness of the transient burst in spiking activity accompanying SWRs and D-U transitions depended on the state of the target region, which varied with sleep depth as operationalized by PSS.

### Modulation of SWR rate by DOWN states is restricted to mouse default mode network

We next asked whether the putative bi-directional interaction observed between hippocampus and RSC extended to other neocortical regions. We first binarized our widefield data into UP and DOWN states using a pixel-wise 25^th^ percentile cut-off, which produced the best alignment of extracellularly and optically detected DOWN states in RSC (**Fig. S8A)**. We then plotted deconvolved widefield activity (**Fig. 6Bi**), RSC MUA (**Fig. 6Bii**), and SWR incidence (**Fig. 6Biii**) surrounding these DOWN states in 7 selected neocortical regions (**Fig. 6A**), spanning medial network (or DMN; red) and somatic sensorimotor networks (blue; networks as determined anatomically in [53]). While DOWN states were reliably detected across these regions (**Fig. 6Bi**; **Fig. S8F**, dotted lines), RSC MUA only followed widefield-detected DOWN states in RSC and regions in mouse medial network, as expected given their dense anatomical connectivity (**Fig6. Bii**; **Fig6. Ci**; **Fig. S8D-F**). Paralleling this, a decrease in SWR rate was observed during DOWN states detected across the medial network (positive SWR modulation index; see Methods; **Fig. 6Cii**), but not somatic sensorimotor networks. This effect was pronounced with longer DOWN state duration (**Fig. S8G**), which occupied greater cortical area.

**Figure 6.**
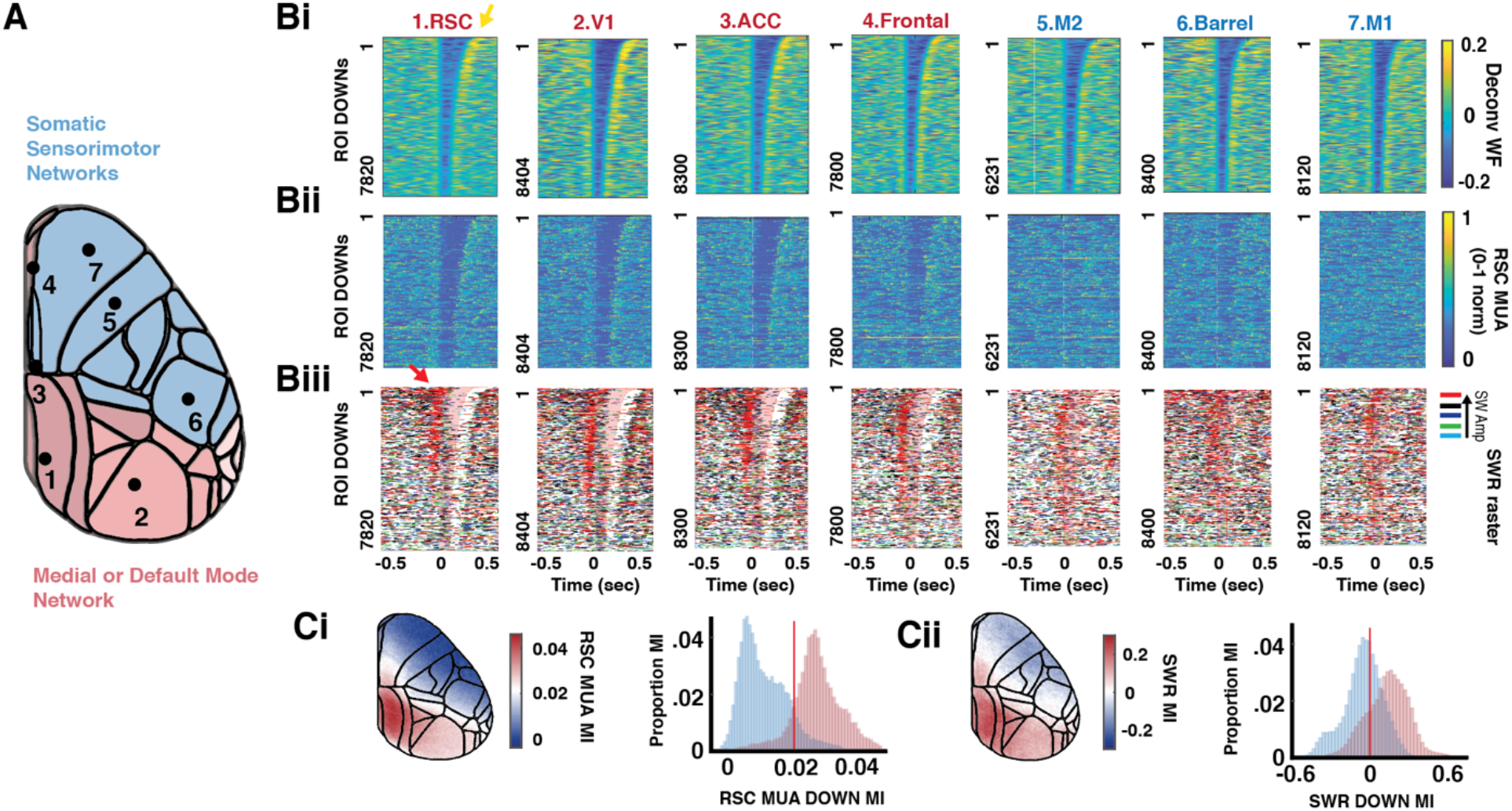
Probability of SWRs surrounding DOWN states across dorsal neocortex. **A**. Map of regions visible in imaging FOV, color-coded by membership in medial network (red) or somatic sensorimotor networks (blue), as in [39]. Numbered regions correspond to columns in Bi-iii. **Bi**. Deconvolved widefield activity surrounding widefield-detected DOWN states in the region specified (25^th^ percentile of pixel WF values and below = DOWN state), as described in **Fig. S8** and **Methods**. Sorted by duration DOWN for an example mouse, separately in each region. **Bii**. RSC MUA surrounding the same DOWN states for each region. **Biii**. Raster plot of SWRs surrounding the same DOWN states, color-coded by SWR amplitude quintiles (small to large: green, cyan, blue, black, red). Note that large amplitude SWRs (red) precede U-D transitions for long DOWN states, red arrow. **Ci**. Average modulation index (MI; see Methods) of RSC MUA by DOWN states detected across all pixels and all mice; positive MI corresponds to higher RSC MUA during UP than DOWN for the given pixel (see Methods for details); *Left*, MI plotted on dorsal map, *Right*, distribution of same values separated by medial (red) and sensorimotor networks (blue). **Cii**. Average modulation of SWRs by DOWN states across all regions; *Left*, MI plotted on dorsal map, *Right*, distribution of same values separated by medial (red) and sensorimotor networks (blue).

To examine DOWN state topography surrounding SWRs, we plotted the average probability of DOWN states surrounding SWR peaks, separated by small and large amplitude SWRs (**Fig. 7Ai, Bi**; t = 0 sec). Consistent with our electrophysiological and optical observations (**Figs. 1, 5**), SWRs were preceded by a significant increase in UP state probability localized to mouse medial network beginning 120 ms before SWR occurrence (**Fig. 7A,B**, red). Whereas small-amplitude SWRs occurred during a DOWN state that remained largely confined to RSC, large-amplitude SWRs occurred during UP states and were followed by DOWN states in RSC and lateral M1/M2 (**Fig. 7Bi**, arrows at 30ms; **Fig. 7Bii**, white outlines) that then spread across neocortex, as measured by a shift in DOWN onset latencies across adjacent cortical regions (**Fig. 7Bii**). DOWN state onset in RSC was followed by DOWN states in visual and somatosensory regions (**Fig. 7Bii**, white to blue outlines). DOWN state onset in M2 and M1 was followed by DOWN states in midline prefrontal, anterior cingulate, and somatosensory regions. DOWN states terminated in V1 and barrel cortex. This suggests large amplitude SWRs are followed by DOWN states initiated in RSC and/or M1/M2 that then invade much of the neocortex with trajectories following cortico-cortical anatomical connectivity (see **Suppl. Movie 4)**.

**Figure 7.**
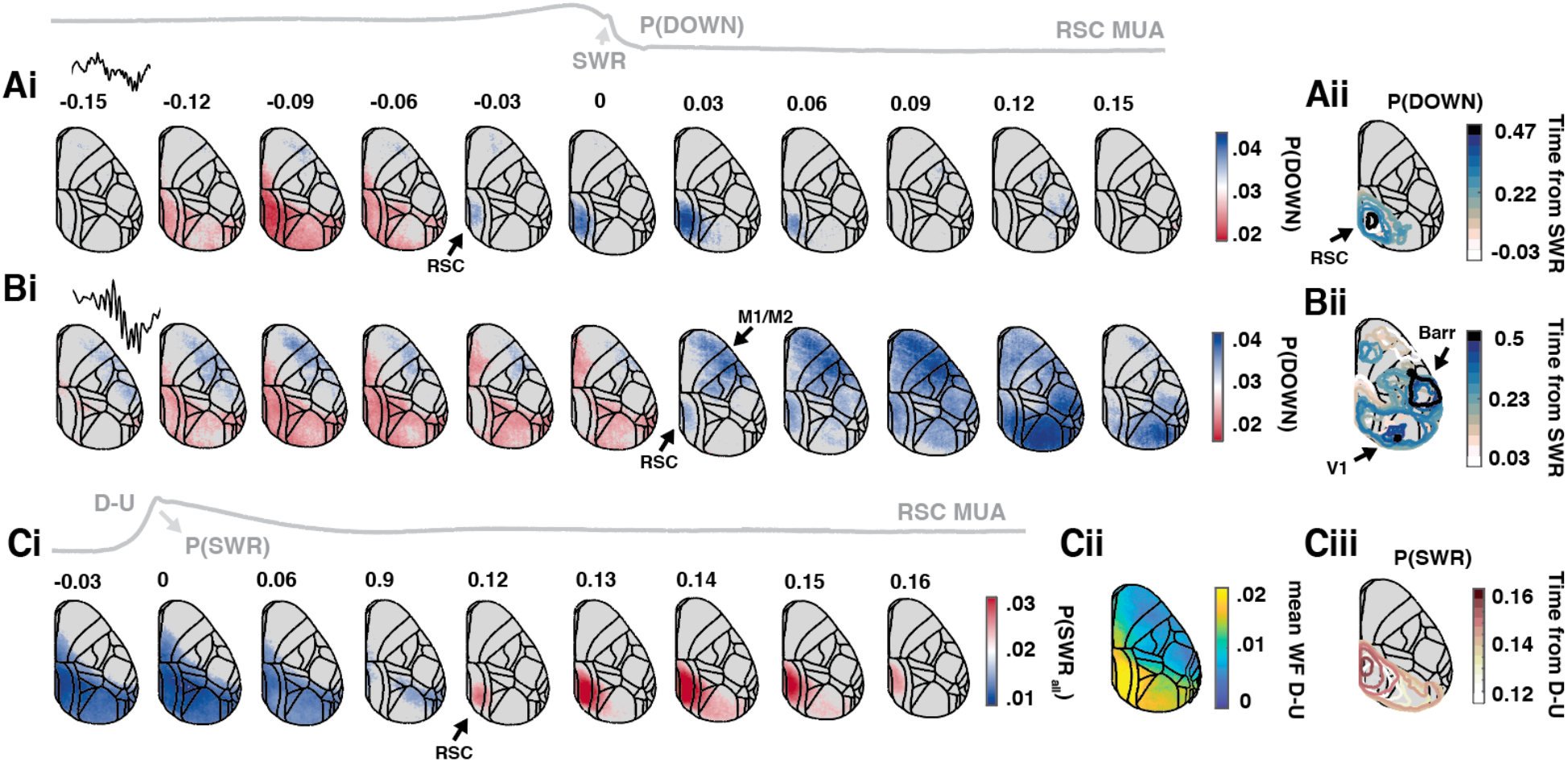
Average topography of putative interaction between hippocampal SWRs and neocortical DOWN states. **Ai**. Average probability of DOWN state occurrence across all pixels aligned to low amplitude SWRs (amplitude quintile 1 of 5; t = 0, peak of SWRs). Colored portion of plots denotes the timepoints at which the given pixel is above (blue) or below (red) a 95^th^ percentile bootstrapped confidence interval, obtained by shuffling SWR peak times across all SWRs and re-computing cross correlograms (n=500). **Aii**. Outline of DOWN states from the onset of DOWN in RSC (white outline) to a sink in RSC (dark blue outline), colored by latency with respect to SWR peak. **Bi**. Same as Ai but for SWR amplitude quintile 5 of 5. Note onset of DOWN states 30 ms following SWR peak in both RSC and regions across sensorimotor network. **Bii**. Outline of DOWN states from onset of DOWN in RSC and sensorimotor regions (white outlines) to sinks in V1 and barrel cortex (dark blue outlines), colored by latency with respect to SWR peak. **Ci**. The probability of SWR occurrence aligned to D-U transitions (t = 0) for every pixel. Colored portion of plots denotes the timepoints at which the given pixel is above (blue) or below (red) a 95^th^ percentile bootstrapped confidence interval, computed as in Ai and Bi but with shuffled D-U transition times. **Cii**. Mean widefield activity within 20 ms of the D-U transition for each pixel. **Ciii**. Outline of significant increase in P(SWR) following D-U transitions for successive frames.

To examine the topography of K-complex impact on hippocampal SWRs, we plotted the average probability of SWRs surrounding the DOWN-UP transition for every pixel (**Fig. 7Ci**; t = 0 sec). A sustained decrease in the probability of SWRs following the D-U transition was observed across the medial network, followed by a peak in SWR probability at ∼120 ms after D-U transitions in RSC that spread toward visual areas, eventually returning to RSC (**Fig 7Ci & iii**; **Supp Movie 5**). Average widefield activity at the D-U transition was greater in the medial network than in somatic sensorimotor networks (**Fig. 7Cii**), paralleling the regions for which SWRs were time-locked to D-U transitions.

### Model of weakly-coupled excitable systems accounts for hippocampal-retrosplenial interactions

We hypothesized that the interactions observed between hippocampal SWRs and RSC DOWN states result from weakly coupled excitable systems [34]. We modeled RSC and HPC each as an adapting inhibition-stabilized network (aISN, **Fig. 8A**, see **Methods**) [54] with slow feedback on excitatory activity [34], [55], corresponding to adaptation in the hippocampus [56], [57] and Ih in the cortex [58]–[60].

**Figure 8.**
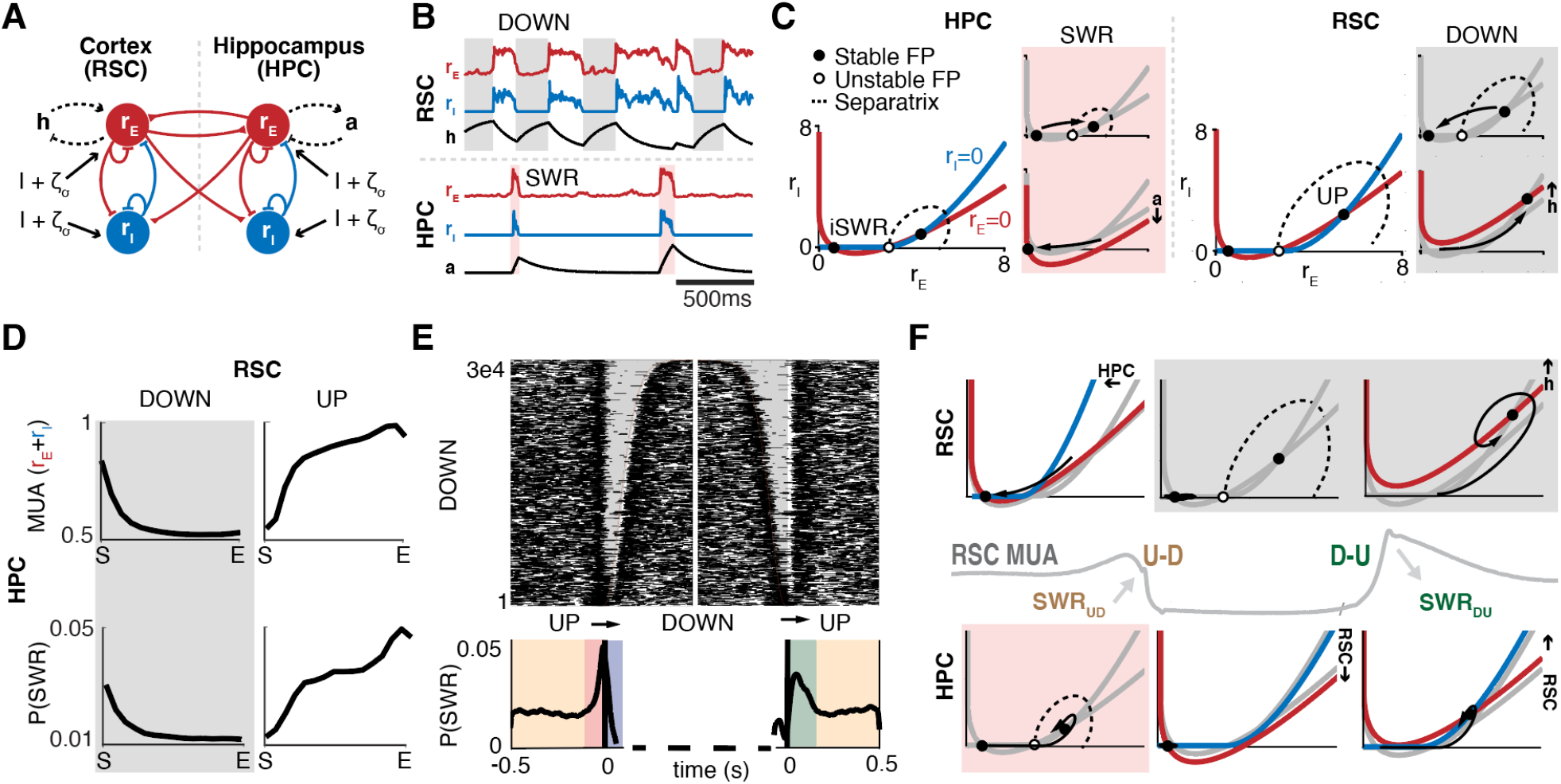
Model of the bidirectional interactions between Hippocampus and Retrosplenial Cortex. **A**. Two-region firing rate model of HPC and RSC with long-range projections between the two regions. Each region comprises of recurrently connected Excitatory (E) and Inhibitory (I) populations with independent background noise. The E populations are subject to a slow feedback current (h-current (h) in RSC, adaptation (a) in HPC, see Methods). **B**. Model simulation outputs for E and I populations in the two regions, and feedback currents. **C**. I-E phase planes for RSC and HPC. Both regions show two stable steady states (a DOWN and an UP state for RSC and an iSWR and a SWR state for HPC). The basin of attraction for each steady state is bounded by a separatrix passing through an unstable fixed point (FP). In the hippocampus (left), a transition from the iSWR to the SWR state engages the adaptative current, which destabilizes the SWR state. In the cortex (right), a transition from the UP to the DOWN state engages the h-current, which destabilizes the DOWN state. **D**. From top to bottom: HPC MUA and P(SWR) plotted as a function of time-normalized RSC UP and preceding DOWN states (compare to **Fig. 3). E. Top**. Raster plot of all SWRs surrounding the DOWN state. Note as in experimental data, clustering of SWRs around UP and DOWN state transitions. **Bottom**. P(SWR) surrounding state transitions reveal a peak before the U-D transition and after the D-U transition (compare to **Fig. 4**). **F**. Analysis of the phase planes for SWR-UP/DOWN interaction. (**i**, SWRUD) Increased hippocampal activity in the SWR state displaces the RSC nullclines, destabilizing the UP state fixed point and pushing the trajectory to a DOWN state. (**ii**) Low RSC activity in the DOWN state lowers the HPC E nullcline, reducing the P(SWR). (**iii**, SWRDU) Activation of the h-current during the DOWN state results in increased RSC activity following the D-U transition. High RSC activity displaces the HPC nullclines, destabilizing the iSWR fixed point and pushing the trajectory to a SWR.

In the presence of noise, the aISN model generates alternation dynamics with asymmetric durations of UP/DOWN states in RSC and SWRs/inter-SWR intervals (iSWR) in HPC ([34], **Fig. 8B, Fig. S9A**), which were used to select model parameters that best matched the data (**Fig. S9A**). These duration statistics emerge because both populations spend their time in complementary excitable states (low-rate iSWR in HPC; high-rate UP in RSC; **Fig. 8C**). In HPC, noise can cause a transition to a transiently stable high-rate SWR state, which is subsequently destabilized by the effect of adaptation (**Fig. 8C**, red shading). In modelled RSC, noise can cause a transition to a transiently stable DOWN state, which is subsequently destabilized by the effect of Ih (**Fig. 8C**, gray shading).

In addition to each region’s local connectivity, we coupled RSC and HPC using excitatory projections that targeted the excitatory and inhibitory populations in the partner region (**Fig. 8A**, Methods). This coupled network exhibited increased incidence of SWRs prior to DOWN states (**Fig. 8E**, compare to **Fig. 4D-E**), decreased hippocampal population rate and pSWR during cortical DOWN states (**Fig. 8D**, compare to **Fig. 3**), and increased pSWR following cortical DOWN-UP transitions (**Fig. 8E**, compare to **Fig. 4D-E**), as in our experimental findings. Analysis of the phase planes revealed that these temporal relationships emerged because the influence of each region on the other modulates the stability of fixed points, and thus the probability of transitions, at critical times (**Fig. 8F, Suppl. Movie 6**). During a SWR, increased drive from the hippocampus decreases the stability of the RSC UP state, increasing the probability of an U-D transition (**Fig. 8Fi**). During the DOWN state, lower drive from RSC decreases HPC firing rate during the hippocampal iSWR and increases its stability, decreasing the probability of an iSWR→SWR transition (**Fig. 8Fii**). Following the DOWN state, Ih transiently increases the firing rate of the CTX UP state fixed point which provides increased drive to HPC, thus decreasing the stability of the hippocampal iSWR state and increasing pSWR (**Fig. 8Fiii**).

Further analysis of the model revealed two additional insights. First, the ability of SWRs to evoke a cortical DOWN state relied primarily on the influence of hippocampal activity on cortical interneurons (**Supp Figure B**), as has been observed experimentally with hippocampo-cortical [61], [62] and cortico-cortical [63] projections. Second, the temporal relationships observed between SWRs and DOWN states relied directly on bi-directional interaction between HPC and RSC, as a “lesion” of CTX->HPC projections resulted in a loss of DOWN state-modulation of hippocampal MUA and thus modulation of pSWR (**Fig. S9C**). Conversely, lesion of the HPC->CTX projection removed the increased probability of SWRs at U-D and D-U transitions (**Fig. S9D**). Together, these results indicate that a mechanism involving coupled “excitable” systems is sufficient to explain the putative state-dependent, bi-directional interaction observed between HPC and RSC.

## DISCUSSION

Using a combination of wide-field imaging of mouse dorsal neocortex and electrophysiological recordings from the RSC and hippocampus, we found evidence of a topographically confined, bi-directional interaction between hippocampus and neocortex, which varied in strength with ultraslow fluctuations in arousal level. In addition to the modulation of SWR rate by UP/DOWN states in the default mode network, population-level state transitions in one structure had a precise temporal relationship with state transitions in the other. From cortex to hippocampus, SWRs followed rebound excitation at D-U transitions, or K-complexes, in the default mode network with a characteristic latency. From hippocampus to cortex, large amplitude SWRs were followed by an increased probability of DOWN states in RSC and antero-lateral motor areas, which spread following cortico-cortical connectivity. A model of weakly-coupled excitable systems accounted for the major experimental observations.

### Putative bidirectional hippocampal-neocortical interaction

Our findings support and extend previous work suggesting a hippocampal-neocortical “dialogue” during NREM sleep. Previous electrophysiological experiments often recorded from the hippocampus and a single partner region. As a result, mechanistic hypotheses proposed based on the observed temporal relationships varied, including that SWRs trigger either UP states or DOWN states, or that the neocortex primes the spike content of SWRs [8]–[16], [64]. Recent imaging experiments attempted to address these contradictions by considering regional variation in coupling, but either lacked the temporal resolution needed to resolve direction of interaction, did not record during NREM sleep, or arrived at hypotheses that differ from ours [31]–[33].

#### From neocortex to hippocampus

Our experiments show that hippocampal spiking activity tracks UP/DOWN states in neocortical regions restricted to mouse default mode network, with the most pronounced covariation between RSC and HPC during deep NREM. Previously referred to as ‘frames’ of co-activity [9], [10], this covariation may be enabled by common third-party drive, for example from subcortical sources [65], [66]. Another possibility is that the traveling UP/DOWN states characteristic of NREM sleep spread to RSC or entorhinal cortex, monosynaptic partners of HPC, which in turn directly drive hippocampal circuits. In support of the latter, in our model, increased input to HPC during cortical UP states increases the excitability of HPC. This caused an increase in both HPC population rate and SWR rate, due to an increase in the ease with which noise or external perturbation can ‘kick’ HPC into a SWR state. In support of this scenario, it was previously reported that both the firing rates of hippocampal neurons and SWR incidence decrease during bilateral optogenetic silencing of the medial entorhinal cortex [13]. The excitability of hippocampal and cortical populations has also been demonstrated to increase with deepening NREM [34], which is reflected in the increased modulation of HPC by RSC UP/DOWN states with deepening sleep.

In addition to the modulation of hippocampal excitability by UP/DOWN states and NREM depth, a disproportionate number of SWRs occurred following DMN D-U transitions at a fixed lag (SWRDU). The putative trigger for SWRDU is the rebound excitation following D-U transitions, known as the K-complex in scalp EEG recordings. Our model supports our interpretation of these observations. In the model, D-U induced k-complexes occur because activation of the h-current during RSC DOWN states results in transient rebound excitation at the D-U transition prior to settling into an UP state. This D-U ‘rebound excitation’ destabilizes the inter-SWR state in the HPC population, thus increasing the probability of SWR occurrence.

Of note, the increase in HPC excitability lagged behind the onset of UP states in RSC and other DMN regions. Mirroring this, SWRDU did not occur in tandem with k-complexes, but rather followed D-U transitions in RSC with a delay of 120 ms (**Suppl. Movie 5B**). An explanation for this delay is not readily captured by our model, even with delayed differential equations (see Methods). It is possible the excitatory drive from RSC is not direct, and occurs primarily via a polysynaptic pathway through either entorhinal cortex or thalamus [13], [67]. However, a similarly long delay has been observed between entorhinal cortical D-U transitions and SWRs [9]. An alternative possibility is that excitatory input drives dentate granule cells, which exert a transient inhibitory effect on CA3 pyramidal cells, via feed-forward inhibition [9], [15], [68], and that the release of those CA3 pyramidal cells from hyperpolarization induces synchronous rebound spiking [69], [70]. Multi-site recordings in RSC, entorhinal cortex, hippocampus, and thalamus, or brief optogenetic hyperpolarization of CA3 neurons, will be needed to test these hypotheses.

#### From hippocampus to neocortex

In the reverse direction, as SWR amplitude and depth of sleep increased, the probability of retrosplenial cortical DOWN states following SWRs at a fixed lag also increased (SWR_UD_; [12], [19], [71]). This temporal relationship is not without precedence, as in humans DOWN states often follow SWRs [72], and interictal epileptiform events in the hippocampus reliably induce DOWN states in both humans and rodents [7]. Our model suggests a mechanism by which SWR-induced DOWN states could occur. A SWR transiently destabilizes the UP state via a strong drive of the local cortical inhibitory population, resulting in increased probability of transition to a DOWN state. Deepening NREM sleep further destabilizes DOWN states [34], contributing to this effect. This mechanism is corroborated by a recent paper that optogenetically stimulated hippocampal terminals in RSC, and found an increase in the firing rate of inhibitory, but not excitatory cells, followed by a DOWN state [73]. In our widefield data, we further observed that sufficiently large amplitude SWRs were followed by DOWN states in RSC or anterolateral motor regions that then spread across much of the neocortex, with average sinks in the barrel and primary visual cortical regions. This may be facilitated by cortico-cortical or thalamo-cortical projections. For example, RSC is a ‘hub’ in the default mode network [74], [75], and shares dense bi-directional projections with regions across the visual hierarchy. SWRUD could ultimately lead to a DOWN state in V1 via induction of a DOWN state in RSC that then propagates along hierarchically connected visual areas. Alternatively, DOWN state induction in early sensory areas could happen via thalamo-cortical disfacilitation, supported by the observation that numerous thalamic nuclei are silenced during SWRs [19], [23], [76], and the larger the amplitude SWR, the more global it is along the longitudinal axis of the hippocampus [19]. Overall, these observations suggest that SWRUD events exert an influence on neocortical activity proportional to SWR amplitude that then propagates across neocortex.

An unexpected observation, in light of previous claims [14], [17], [18], was the absence of SWRs preceding and thus putatively inducing UP states. We observed only a small fraction of SWRs during DOWN states, often timed by the K-complex of a preceding short-duration UP state at ∼120 ms. The failure of SWRD to induce a D-U transition could be explained by their low probability, low amplitude, or refractoriness of the target circuits. In line with this latter explanation, SWRs during DOWN states evoked EPSPs in entorhinal neurons but failed to discharge them [9], preventing the propagation of excitation. We also note that there were more U-D and D-U transitions than SWRUD and SWRDU events, implying that only a fraction of these transitions were induced by or induced a SWR. One possible explanation for this is that traveling slow oscillations [26] observed in DMN or RSC may fail to invade the entorhinal cortex, the primary input to the hippocampus. Another explanation, afforded by our model, is that both regions are only weakly coupled, and thus capable of noise-driven transitions independently of one another.

### Putative functions of SWR types

The ability to distinguish SWRs by their timing with respect to neocortical UP and DOWN transitions could help disentangle the direction of spike transmission between hippocampus and neocortex, and thus the mechanistic contribution of these ‘SWR types’ to memory. One possibility is that the observed SWR types support distinct functions, such as encoding, consolidation, or priming of recalled events. In a recent study, hippocampal reactivation occurred during prefrontal cortical UP states, whereas the strongest coordination between RSC and hippocampus occurred during U-D transitions in RSC [77]. SWRD events, some of which may be triggered by K-complexes, can sporadically activate a few neocortical pyramidal cells during the DOWN state. This sporadic spiking during DOWN states has been suggested to be the critical driver of consolidation of recently acquired experience [78]. However, this explanation alone would leave the function of the great majority of SWRs unexplained, other than serving subcortical, autonomic functions [79]. In contrast, another study emphasized the importance of distinct brain-wide coordinated and uncoordinated SWR events during UP states [80].

A complementary hypothesis is that the four types of SWRs are better understood as part of a multi-regional ‘dynamical motif’ enabling systems consolidation [81], facilitated by the excitable regimes characteristic of NREM sleep [34]. SWRs, if sufficiently large, may induce a DOWN state (SWRUD). This DOWN state may invade thalamus, inducing a thalamo-cortical spindle [82], and the rebound excitation from the D-U transition may then initiate a SWR burst in HPC that is coordinated with that induced spindle. In support of this, memory reactivations in humans occur when SWRs are coupled to slow oscillations and spindles but not during solitary slow oscillations or spindles [83]. Further, SWR bursts are likely important for consolidation in light of reports that long-duration neuronal spike sequences, reflecting long trajectories in a previously experienced environment, span several hundred milliseconds and often abridge two or more SWR events occurring in a burst [84]. Whereas SWRDU are more likely to reflect burst onsets, SWRUD may play a role in ending both a SWR burst in HPC and an UP state in CTX. One can speculate that the ensuing silence serves as the truncation of coordinated exploration along a given attractor, or expression of a memory trace, allowing exploration of the next [24].

### Arousal levels affect interregional perturbation

Ultraslow and infraslow fluctuations in arousal level have long been observed in both humans and rodents [20]. However, the link between these slow timescale changes and fast timescale hippocampal-neocortical interaction has remained elusive, resulting in largely separate rodent and human literatures. We suggest that the dynamical regime, and thus excitability, of brain circuits fluctuates across ultraslow and infraslow timescales, likely due to the slow changes in neuromodulatory tone accompanying transitions in arousal level [85], [86]. Ultraslow fluctuations reflect global changes in arousal level, whereas infraslow fluctuations reflect changes in regime within resting state networks. Given the hypothesized fluctuations in regime, these slow rhythms reflect the propensity with which the regions belonging to the given resting state network can be perturbed [34]. For example, an ‘active’ DMN corresponds to an increased rate of SWRs and DOWN states in DMN regions [27] which arise due to the more ‘excitable’ regime the DMN is in, facilitating inter-regional communication within but not across resting state networks. Finally, SWR ‘types’ arise because of the transition from less to more excitable regimes over the course of deepening sleep. If sufficiently excitable or if the perturbation is sufficiently large, SWRs (SWRUD) can cause DOWN states, and D-U transitions can cause SWRs (SWRDU). These perturbations can then propagate as a function of the state and anatomical connectivity of the downstream structure. This provides a mechanism by which SWR perturbation can propagate along the neocortical hierarchy, mediated by sleep depth.

We did not distinguish explicitly between wake and sleep SWRs. This may be considered a caveat, given the distinct functions they are often assigned [85], [87], [88]. However, our observations and previous results [89] do not support a clear delineation between wake and sleep, but rather a transition toward an increasingly ‘excitable’ neural regime as an animal moves through quiet wake to deep NREM sleep states. Supporting this notion, UP-DOWN states are present during quiet wake, but are notably more localized, as is the impact of perturbation via SWRs [15], [89]. Further experiments are needed to reveal whether waking and NREM SWRs are qualitatively different in their interaction with neocortex, or whether they are better understood as existing along a continuum.

## Acknowledgments

We thank members of the Basu and Buzsáki lab members for support and feedback. This work was supported by NIH grants MH122391 (GB and JB), U19 NS107616 (G.B.), R01 NS109362 (JB), R01 NS109994 (JB), R01MH062346 (X.J.W.), and R90DA060339 (E.C.).

## Author contributions

RS, GB and JB designed the research. RS, NM, and MV performed the research. RS analyzed the data. EC modeled the data guided by RS, DL, and XJ. RS, GB and JB wrote the paper with contribution of all authors.

## Additional information

Extended data is available for this paper at https://doi.org/… (Supplementary Material)

## Supplementary information

The online version contains supplementary material available at https://doi.org/….

## Data availability

The data of this study are publicly available on the Buzsaki Lab web page (https://buzsakilab.com/wp/resources/).

## Code availability

The code used for this study was adapted from the buzcode repository (https://github.com/buzsakilab/buzcode).

