## SupplementalMaterial for "Topography of putative bidirectional interaction between hippocampal sharp wave ripples and neocortical slow oscillations"

#### SUPPLEMENTAL MATERIALS

#### EXPERIMENTAL METHODS

##### *Animal handling*

All experimental procedures were conducted in accordance with the National Institutes of Health guidelines and with the approval of the *New York University Grossman School of Medicine Institutional Animal Care and Use Committee (IACUC)*. C57BL/6J-Tg(Thy1-GCaMP6f) GP5.17Dkim/J (Jackson laboratories, Stock #:**025393**;(Dana et al., 2014)) mice were used for all data collected; all males. Mice were kept in the vivarium on a standard 12-hour light/dark cycle and were housed 2-5 per cage before surgery and then alone following the surgery. Mice were provided food and water ad libitum throughout experiments, and experiments were conducted at the beginning of the light cycle to maximize sleep duration and quality.

##### *Surgical procedures*

Mice were anesthetized with isoflurane, homeothermically maintained, and monitored using pulse oximetry. In a fully aseptic environment, the scalp over the craniotomy area was resected. Skull thinning was then performed with constant saline irrigation, avoiding excessive drilling of the sutures but ensuring the removal of the outer and spongy bone layer. The thinning area extended over the entire dorsal right hemisphere, from the frontal cortex to the posterior visual cortex. Once drilled to translucency with only the inner compact bone layer remaining, the dry surface of the skull was coated with a thin layer of cyanoacrylate (gel-type Loctite Super Glue), which serves to index-match the rough surface of the skull, preventing bone re-growth and provide mechanical protection. The skin around the thinned skull preparation was sealed using cyanoacrylate tissue adhesive (e.g., 3MTM VetbondTM Tissue Adhesive). A thin coating of clear nail polish was then applied on top of the cyanoacrylate to further index-match the skull's surface and reduce scatter. Finally, a ground screw coupled with a 0.005” stainless steel wire (A-M Systems, #792800) was implanted in the skull above the cerebellum, and a custom 3d-printed head post was fixed to the surface of the skull using dental cement. Mice were then allowed to recover for one week and were inspected for response to visual stimuli to confirm the expected expression and dynamics of GCaMP6f.

In a second surgery, performed 5 days to one week following the first surgery, mice were implanted chronically with a 64-site or 128-site linear silicon probe (H3; Cambridge Neurotech) at a 60$^{o}$ angle, mounted onto a Cambridge Neurotech Nanodrive. The probe spanned deep (64-cite) or deep and superficial (128-cite) layers of retrosplenial cortex and area CA1 of the hippocampus ipsilateral to the imaging field of view, after being lowered from a craniotomy at 0.5 mm DV and 2.95 mm AP coordinates in the contralateral (left) hemisphere (Fig. 1). The probe and nanodrive were then enclosed in the remaining components of the 3d printed head-post.

##### *Behavior*

After recovery for 1 week, animals were gradually habituated to head fixation (modified rivets system; Osborne & Dudman, 2014) for 5 to 7 days, where the amount of time spent head-fixed increased from 10 minutes to two hours. During both habituation and head-fixed experimental sessions, animals were free to run or rest on a 1.9-m length custom treadmill, during which no reward was delivered. Animal behavior was monitored during head-fixation using a small camera (Basler ace acA1300-60gm-NIR GigE Camera) under IR illumination, and a hall effects sensor (Littelfuse 55140-3H-02-A) placed at the axel of one treadmill wheel. Mice were encouraged to sleep using the following strategy: performing imaging sessions at the transition to their sleep cycle (i.e., daytime), increasing the force needed to move the treadmill but still allowing movement, and using a heated treadmill platform that partially enclosed their bodies. Home-cage sleep sessions allowed the mice to behave freely while plugged into the recording cable. Both head-fixed and home-cage setups were Faraday-protected using a metal mesh connected to the ground.

##### *Electrophysiological Recordings*

Electrophysiological recordings were conducted using an Intan RHD2000 interface board with sampling at 20 kHz (IntanTech). Signals were recorded in a unipolar manner against a reference and common ground (cerebellar screw) and digitized by a head-mounted preamplifier. Frame times for the behavior-recording camera and the widefield imaging camera signal were recorded as digital inputs to the Intan system. Probes were gradually lowered until sharp wave ripple (SWR) depth profiles were clearly visible and then left in place for the duration of the experiment.

##### *Widefield Recordings*

Widefield imaging was performed on the right hemisphere of head-fixed mice through the thinned skull preparation using the MVX-10 Macroscope (Olympus) during simultaneous chronic electrophysiological recordings of ipsilateral retrosplenial cortex and CA1. Data were recorded using an Andor Zyla sCMOS camera with dual-wavelength LED imaging (cool LED PE-4000 system) and camera control by NIS Elements software (Nikon). 16-bit images were acquired at a rate of between 66.66 Hz and 100 Hz using global shutter mode. Alternating illumination between 470 nm and 525 nm on a per-frame basis was used in some cases, and illumination with a third wavelength at 630 nm was used for a small subset of videos. Illumination was collected via a 500-nm long pass emissions filter.

##### *Histology*

After experiments, mice were anesthetized with an injection of 150 mg/kg ketamine and 10 mg/kg xylazine. After checking for the absence of reflexes, animals were transcardially perfused by inserting a 27-gauge needle into the left ventricle of the heart while simultaneously severing the right atrium. Following perfusion with 1X PBS, 4% paraformaldehyde in PBS was circulated in the animal. The brain was removed and fixed overnight in 4% paraformaldehyde at 4ºC and then sectioned into 60-µm thick horizontal slices using a microtome (Leica VT1000S) after washing to undergo immunohistochemistry. The slices were mounted on imaging slides in Vectashield Hard Set Mounting Medium with DAPI (Vector Laboratories). Brains were then imaged under a florescent microscope under 350 nm to capture DAPI and 470 nm wavelengths to capture the GCaMP signal.

#### DATA ANALYSIS

Data analysis was performed using custom software written in Matlab (MathWorks) and adapted from the Buzsaki Lab code repository (https://github.com/buzsakilab/buzcode).

##### *Electrophysiological data preprocessing*

Data files for paired head fixed and homecage recording sessions were concatenated and downsampled to 1250 Hz for LFP analysis.

##### *Widefield Data preprocessing (adapted from* **Peters et al., 2021)**

All wide-field data were de-noised via compression using singular value decomposition of the form F = USV. The input to the SVD algorithm was F, the pixels x time matrix of fluorescence values. The outputs were U, the pixels x components matrix of template images; V, the time x components matrix of component time courses; and S, the diagonal matrix of singular values. The top 200 components were retained, found to be inclusive of a threshold determined by the Marchenko-Pastur distribution of a random matrix (Veraart & Novikov, 2016); a more theoretically-motivated noise threshold, but ultimately more time extensive and unnecessary to compute given sufficient compression with 200 components. All orthogonally invariant operations (such as deconvolution, event-triggered averaging and ridge regression to predict cortical activity from the wide-field signal) were carried out directly on the matrix **S*V**, allowing a substantial saving of time and memory.

Hemodynamic effects on fluorescence were removed as in (Ma et al., 2016; Peters et al., 2021), by regressing out a green 525 nm widefield channel, reflecting changes in total blood volume, from the calcium-dependent signal obtained with blue illumination (470 nm). To achieve this, both signals were band-pass filtered in the range 7–13 Hz (heartbeat frequency, expected to have the largest hemodynamic effect). Pixel traces for blue illumination (470 nm) were then temporally resampled to be concurrent with green illumination (525 nm, as colors were alternated), and a scaling factor for the given pixel-wise regression was fit across colors for each pixel. The scaled fluorescence traces from 525 nm green light illumination were then subtracted from the fluorescence traces emitted from 470 nm blue light illumination. To correct for slow drift, hemodynamic-corrected fluorescence was then high-pass filtered over 0.01 Hz and $\Delta F/F0$-normalized by dividing by the average fluorescence at each pixel within the session.

Wide-field videos were then aligned across sessions by rigid registration of the average green-illuminated image to the Allen Common Coordinate Framework52 (Wang et al., 2020) using two anatomical coordinates, lambda and bregma (marked during surgery). In a subset of mice, visual stimuli were presented and average deconvolved activity was confirmed to show a response in

V1 (see supplemental figures). In all mice, the closest correlation with RSC population rate was in RSC, as expected.

In some cases, for very long videos, SVD was performed on segments of individual videos. When this was done, U matrices differed across sub-videos, leading to data reconstruction occurring across distinct principal axes. To combat any potential issues arising from this, all U matrices (200 components) were concatenated, and SVD was performed on this matrix; data were then reconstructed using this 'master' U matrix.

Fluorescence was deconvolved using a kernel fit from predicting retrosplenial cortical (RSC) multiunit activity from wide-field GCaMP6f fluorescence using ridge regression. This kernel was estimated by using simultaneous wide-field imaging and electrophysiological recordings in RSC. The single pixel with the highest correlation to the population rate before deconvolution was used for this purpose, and the population rate was calculated by binning spiking data for a spike-sorted session per mouse. 10-fold cross-validation was performed, and a lambda of 0.02 was used for regularization. The final deconvolution kernel was a mean of maximum-normalized kernels across recordings divided by the sum of squared weights across time. The deconvolution kernel was biphasic and roughly similar to a derivative filter (that is, [−1,1]), consistent with rises in the GCaMP signal corresponding to periods of spiking, as in (Peters et al., 2021).

Analysis of traveling waves was performed using the Optical Flow Connectivity Toolbox, (Afrashteh et al., 2017). Note that vector fields were computed using average SWR-triggered videos and thus denote changes in local widefield values from frame to frame on average, which may or may not be representative of individual trajectories. For our purposes, this is sufficient to give a sense of changes in activity over time on average.

##### *Multiple unit activity (MUA) extraction*

MUA extraction was implemented by MUAfromDat.mat (git: buzcode) via the following steps: First, a user-specified channel from the raw .dat file in the region of interest was band-pass filtered in the 500 to 5000 Hz frequency range. Then the estimated EMG from the LFP (Schomburg et al., 2014) was used to replace EMG-related artifacts with 'NaN' values. MUA was then normalized between 0 and 1 within each session. Two common methods of MUA extraction, MUA via band-pass filtering vs MUA via the pooling of spike-sorted units, were indistinguishable.

##### *UP/DOWN state detection*

DOWN states were detected using DetectSlowWavesMUA.mat (git: buzcode), parameters: smoothwin = 0.3, startbins = 40, refineDipEstimate = true. Briefly, MUA is detected as described above. If the distribution of the log-transformed MUA values are significantly bimodal (Hartigan's dip test), the trough of this bimodal distribution is identified (bz_BimodalThresh.mat), and anything above this threshold is regarded an UP state; below a DOWN state. A Schmidt trigger (threshold), which uses the halfway points between trough and upper peaks for onset and offset of a given state, was compared to a hard threshold, and no significant difference was found, as expected given the high data quality (Schmidt triggers are a useful control in noisy data).

***Ripple detection and analysis***

Ripples were detected as described previously (Tingley & Buzsáki, 2020), using bz_FindRipples (git: buzcode). Briefly, the raw LFP (1250 Hz) was filtered (130-200 Hz; Butterworth; order = 3) and was transformed to a normalized squared signal (NSS). This signal was used to identify peaks beyond 5 standard deviations above the mean NSS. The beginning/end cutoff of the ripple was defined by a threshold of 2 standard deviations above the mean NSS. Ripple duration limits were between 15ms and 250ms. In addition, estimated EMG from the wide-band recording (Schomburg et al., 2014) was used to exclude EMG-related artifacts. The peak of the ripple (max power value > 5 standard deviations) was defined as time 0 for the ripple. Ripples were defined across the entire duration of a day, including both periods in the maze and the homecage. Ripple frequency was computed as the derivative of the ripple-band filtered LFP during ripple intervals found above. Ripple power is the power in the ripple band found via wavelet decomposition normalized by power in the ripple band during inter-ripple intervals. Sharp wave (SW) magnitude was the log10 normalized value of the sink after calculating the CSD across channels spanning the pyramidal cell layer using the inverse CSD method, which is based on the inversion of the electrostatic forward solution (Pettersen et al., 2006). All SWRs were manually inspected, and any clearly resembling noise was excluded from the analysis.

***Tonic MUA***

Tonic MUA was computed by calculating the median deep RSC MUA across the duration of a given UP state. Each UP state has a single tonic MUA value.

***Average variables across time normalized UP/DOWN states***

Time normalized UP and DOWN states were computed by taking the median value of interest (RSC MUA, HPC MUA, P(SWR), amplitude SWR, P(burst onset SWR)) within 15 evenly spaced bins within either all UP states, or all DOWN states, and then averaging the value of interest across all time normalized UP states or time normalized DOWN states. Because number of bins was kept constant to allow averaging of variable duration UP and DOWN stats, bin time duration was variable.

***Brain State Scoring***

A previously described semi-automated sleep scoring algorithm was used (Watson et al., 2016). It calculates a spectrogram from the raw LFP (1.25kHz) using a 10-s window FFT, sliding at 1 s intervals, at logarithmically spaced frequencies from 1 to 100 Hz. Briefly, this algorithm uses a set of heuristics when examining the EMG, theta band ratio (4-9 Hz divided by 2-16 Hz), and broadband LFP. The estimated EMG is the summed pairwise zero-lag correlation between non-neighboring electrodes (separate shanks; > 200um distance) using the bandpass filtered (300-600 Hz; 3rd order Butterworth) local field potential (Schomburg et al., 2014). The theta band ratio is the 4-9 Hz power from the spectrogram divided by the 2-16 Hz power. The broadband LFP spectrogram was then compressed using principal components analysis (PC1 always corresponding to < 20Hz power). The algorithm then uses these data in a sequence of separations, finding troughs that maximally split peaks in each distribution. In all recordings, manual inspection of scoring was conducted. In some cases, manual curation of algorithm parameters or specific segments of recordings was conducted to identify brain state best.

***PSS Power spectrum slope (PSS)***

We used the power spectrum slope as an estimate for brain state and as a potential index of cortical E/I balance (Gao et al., 2017). For each session, we selected an infragranular cortical channel and extracted the slope of the power spectrum between 4-100 Hz in a 2 seconds interval and a 50 ms sliding window. PSS describes state changes as a single value. Its relationship to classically defined brain state changes can be calibrated and PSS values can then be assigned to these discrete (arbitrary threshold-separated) states. While Brain State scoring provides isolated clouds for active waking (locomotion) and REM sleep, the relationship between quiet awake immobility and NREM sleep is more of a continuum than bimodally separable states (Yamabe et al., 2019).

***Computing SWR phase-relationship with infraslow blood flow signal***

Every neocortical pixel in the 520 nm wavelength (green) channel was filtered in the infraslow frequency range, [.04 to .5] Hz. The phase angles for each timestamp when a SWR peak occurred within NREM epochs were then calculated separately for each pixel using the real component of the Hilbert transform and plotted in a histogram. The circular mean and resultant vector were then calculated using these histograms for each widefield pixel (Berens, 2009). For a null distribution, widefield infraslow phase angles were circularly shifted by a random offset and the phase angle histograms were then recalculated (10 iterations). Resultant p-values are calculated based on this bootstrapped null distribution and plotted for each pixel.

***Spectrograms x infraslow phase***

We first concatenated all NREM epochs for a given session within mouse of a select LFP channel, either from deep-layer RSC or the CA1 pyramidal cell layer. We then calculated a time-resolved wavelet spectrogram by convolving this LFP time series with a family of complex Morlet wavelets (Torrence & Compo, 1998), log2-spaced between 1 and 200 Hz with variable time resolution, allowing for fixed five cycles per frequency. The wavelet spectrogram was then mean-normalized within frequency. Next, we plotted the average power spectrum by infraslow phase (Fig. 2I) by assigning an instantaneous phase of total blood volume (520 nm channel) filtered in the infraslow frequency range (0.04 to 0.5 Hz), to each time-point in the spectrogram, sorting the spectrogram into 20 evenly spaced phase-bins, and then averaging across time within those bins. This allowed us to visualize changes in the power spectrum for the given LFP by phase infraslow (see bz_LFPSpecToExternalVar).

***Identification of SWR types***

In main Figure 4, we observe asymmetric clustering of SWRs around U-D and D-U transitions and ultimately identify four SWR "types" based on their proximity to the nearest U-D and D-U transition: $SWR_{U}$, $SWR_{UD}$, $SWR_{D}$, and $SWR_{DU}$. We note that the $SWR_{UD}$ and $SWR_{DU}$ are partially overlapping categories, as a SWR can both follow a D-U transition at a specific latency and occur just before a U-D transition. For the purpose of our study, grouping SWRs into overlapping categories is not problematic, as we are ultimately testing the hypothesis of bidirectional interaction between regions and do not group SWRs by "type" for the analyses in main Figure 5,6, or 7. We elaborate on our classification of SWR types in **Figure S6.** First, we plotted the distribution of the latency of the specified SWRs from four time points: the latency of all UP SWRs to the closest previous U-D transition (**Fig. S6B**), the latency of all DOWN SWRs following the closest previous U-D transition (**Fig. S6C**), the latency of all DOWN SWRs to the next D-U transition (**Fig. S6D**), and the latency of all UP SWRs from the previous DOWN-UP transition (**Fig. S6E**). From these distributions, latency cut-offs were determined based on peaks in the latency distributions, latency intervals noted on plots, the colors in **Fig. S6B-E** corresponding to the outlined peaks in P(SWR) in **Fig. S6A**. $SWR_{UD}$ in category 'A' (red) occurred within 40 ms of the U-D transition, $SWR_{DU}$ (green) in category 'E' occurred between 50 and 180 ms post-D-U transition, $SWR_U$ in category 'F' (yellow) occurred during an UP state between 180 to 200 ms following a D-U transition, and $SWR_D$ in category 'B' (magenta) occurred between 1 to 80 ms following the U-D transition.

In **Fig. S6F-G,** we assign every SWR to one of 9 non-overlapping categories defined by their proximity to the closest U-D and D-U transitions (numbered 1-9), specified by the combinations of latency intervals below each plot (more generally – how far is a SWR from the nearest U-D and D-U transition, and how long is the UP state within which it occurs). Categories 1-7 occur during the UP state, and 8-9 during the DOWN state. Categories 1-3 comprise $SWR_{UD}$, category 5 $SWR_{DU}$, category 8 $SWR_D$, and categories 6 and 7 $SWR_U$. In **Fig. S6H**, we present SWR raster plots of each SWR type surrounding the D-U transition, colored by the amplitude of the SWRs. We note the four major SWR types whose potential functions we discuss in the manuscript.

***Calculation of DOWN states in widefield data***

Deconvolved widefield data was binarized into putative UP and DOWN states by thresholding each pixel by its 25$^{th}$ percentile value, above which was an UP state and below which a DOWN state. The 25$^{th}$ percentile was chosen because it maximized the KS-distance between RSC MUA values during RSC widefield detected UP vs DOWN states (**Fig. S8A)**. In more detail, we detected putative DOWN states in RSC widefield time series using thresholds evenly spaced between 15 and 40$^{th}$ percentiles, leading to variation in identified DOWN onsets and offsets. We then leveraged our simultaneous RSC widefield and electrophysiological recordings by computing the distribution of RSC MUA values in each of the widefield-determined putative UP and DOWN states (pooled across all videos within a given mouse to control for variation in depth of NREM). The value with the greatest distance between detected UP and DOWN states was the 25$^{th}$ percentile threshold. We then calculated the 25$^{th}$ percentile value for each pixel within each mouse, and applied this threshold to that pixel's time series to identify UP and DOWN states. Although we could only perform the above electrophysiologically-determined widefield threshold within RSC, we found no systematic bias in computed 25$^{th}$ percentile values across regions (**Fig. S8B,C**), as expected assuming an approximately uniform imaging quality.

Given the lower temporal (and thus spatial) resolution of this datatype, we performed an additional quality check, wherein we evaluated changes in DOWN detection quality (operationalized as UP vs DOWN KS-distance) as a function of DOWN duration deciles for

RSC and M1 ROIs (**Fig. S8D-F)**. In short, for the specified ROIs (**Fig. S8F**), DOWN states were split into duration deciles, from short to long. Then, two KS-values were computed (dotted and solid lines in **Fig. S8E**), addressing two distinct questions. First, we asked whether the widefield DOWN detection quality is similar across regions as DOWN state duration varies (dotted lines). It is possible that although long, and thus global, DOWN states are easily detected across regions, shorter duration (and thus local) DOWN states may be missed in regions not central to our investigation. This could be problematic for analyses that vary the duration of DOWN states, such as computing SWR modulation indices. This control analysis was thus done by computing the KS-distance between distributions of UP and DOWN widefield values, for the given pixel, at each duration DOWN decile. Across regions, KS-distances remained similar regardless of duration DOWN – suggesting minimal variation in detection quality (**Fig. S8E,F, dotted lines**).

A second question can be addressed given our concurrent electrophysiological and widefield recordings in RSC - does widefield DOWN detection quality vary with DOWN duration if using RSC MUA. Ideally, when calculating KS-distances using RSC widefield-detected DOWN states to bin RSC MUA into 'UP' and 'DOWN' distributions, we would see minimal change in KS-distance, as is true when using electrophysiologically-detected DOWN states to bin RSC MUA (**S8F, black line**). However, we instead observed an increase in KS-distance with increasing DOWN duration, meaning a systematic improvement in detection quality. This is unsurprising, as it reflects the temporal resolution of widefield data. Fortunately, for our purposes, we exclude DOWN states shorter than 80 ms in duration in order to avoid including other silent events (e.g., 4Hz or spindles) in our results. Overall, even if short-duration DOWN states are less effectively resolved, comparisons across regions remain sound given the equivalent quality of DOWN state detection across regions using widefield time series.

***DOWN state modulation index***

For every pixel, DOWN states detected for the given pixel were divided into duration quintiles, from short to long. For each quintile, a DOWN state modulation index was computed by dividing the total number of SWRs observed during UP states minus DOWN states, divided by the sum of SWRs in those same UP states and DOWN states. To compensate for the variable duration of UP and DOWN states, SWRs were summed only within a 70ms window from the end of each UP state, and end of each DOWN state. (Note: DOWN states with less than 80ms duration were excluded to avoid mid-identification of spindle troughs). Modulation indices were averaged across mice, resulting in one modulation index per pixel (**Fig. 6C**), or one modulation index per pixel per DOWN duration quintile (**Fig. S6G**), plotted in a map in **Fig. 6E**.

***Cross-correlograms of widefield data***

The probability of DOWN states surrounding SWRs (Fig. 7Ai,Bi) and SWRs surrounding U-D and D-U transitions (Fig. 7Ci) was quantified by calculating the cross-correlation between the trigger of interest for a given pixel, SWR peaks sorted by amplitude and D-U transitions respectively, pooled across all mice. This provided a unique map in time, showing the probability of DOWN state transitions across regions surrounding SWRs (Fig. 7Ai) or probability SWRs in a single hippocampal location with respect to U-D and D-U transitions across all regions at fixed lags from those transitions (Fig. 7Ci). The frames in **Fig. 7Ai, Bi, Ci** are still pictures taken from the videos in **Supp. Movies 4 and 5**. Data in Fig. 7 are thresholded by a 95th percentile bootstrapped confidence interval for each pixel, constructed by re-computing the cross-correlograms of interest n=500 times.

***Computing cross-correlograms to test whether the strength of input impacts population-level state transition in downstream region***

We tested the hypothesis that the hippocampus and RSC can bi-directionally "kick" one another by computing cross-correlograms between putative input variables (in the hippocampus to RSC direction: SWRs; in the RSC to hippocampus direction: D-U transitions) and putative response variables (hippocampus to RSC: DOWN states; RSC to hippocampus: SWRs), by reasoning that the larger the input variable, the greater the probability of evoking a state transition in the downstream region. We thus split the input into octiles that varied by input "strength", operationalized as the amplitude of SWRs or magnitude of synchrony at the D-U transition, for hippocampus and RSC, respectively. For each cross-correlogram (CCG) between input times of a given input strength octile and all response times, surrogate data sets were constructed by randomly and independently jittering timestamps on a uniform interval of [-20, 20] ms 1000 times. 99% confidence intervals were then calculated for each time bin, plotted as shaded regions along the CCGs in **Fig. 5B,E** (Fujisawa et al., 2008). The larger the SWR octile, the higher the probability of a DOWN state following that SWR octile at a fixed lag of 30ms±15 ms. The larger the synchrony, the higher the probability of a SWR following the transition at a fixed lag of 120ms±15 ms. For all CCGs in **Fig 5**, time series were combined across animals and variations in input strength were denoted with deepening color (light pink to dark red; or light green to dark green). The same conclusions can be drawn from cross-correlograms of individual mice, suggesting strong evidence of short-timescale bi-directional interaction between hippocampus and RSC via coordinated state transitions.

***Testing the effect of input strength and state of downstream region on probability of evoking a population-level state transition via repeated measures ANOVA***

We next tested for the effect of two independent variables and their interaction on the probability of evoking an RSC DOWN state. For this analysis, we binned SWRs not just by input strength octiles, but also by the arousal level of the animal, taken to be PSS value when a given trigger, a SWR or D-U transition, occurred. Given the fixed lag at which DOWN states occur in response to increasingly large SWRs (30±15 ms), we computed CCGs for each condition, computed a two-way repeated measures ANOVA with values of each CCG at this time bin, and tested for a significant effect of amplitude SWR, state, and their interaction at the session level across mice. As expected, we found a significant effect of both factors and their interaction; the higher the PSS value (corresponding to deeper NREM sleep) and larger the SWR, the higher the probability of an evoked DOWN state. These effects were lost at a control time lag of 200 ms. We followed this same logic in the RSC to hippocampus direction and found a significant effect of both magnitude synchrony, PSS value at the time of the trigger, and their interaction in probability of evoking a SWR at a lag of 120 ms (two-way repeated measures ANOVA). At a control lag of 350 ms, we found a significant effect of local state but not strength input on probability of SWR occurrence.

***Predicting duration DOWN state or magnitude of sharp wave using input strength and local state as predictors in a generalized linear model***

In events where a DOWN state followed a SWR ($SWR_{UD}$), we tested whether the duration of the DOWN state varied with state, in this case, PSS, and input strength (i.e., SWR amplitude) using a generalized linear model with both variables (PSS and SWR amplitude) and their interaction as predictors, and duration DOWN state as the response variable. In this case, we did not bin PSS or SWR amplitude and only performed this binning with average duration DOWN for visualization of results. We found a significant effect of both predictors and their interaction. When we took all SWRs during the UP state and repeated this process for the following DOWN state (rather than just SWRs with a putative causal impact on cortical DOWN state given their occurrence just prior), amplitude SWR was no longer predictive of duration DOWN, further supporting our causal hypothesis. Note, however, that we only found a significant effect of state using PSS, reflecting global arousal level, rather than local state.

In those cases where a SWR followed a transition from a D-U state, we tested whether the magnitude of the sharp wave, reflecting putative degree of input, depended on the strength of the input and state, by using synchrony at the D-U transition, PSS, and their interaction as predictors using the same GLM framework. We found a significant effect of state, but not of strength input or the interaction between the two predictor variables. This suggests that magnitude SW is determined largely by local excitability, which is presumed to change with depth sleep, rather than strength input.

***Statistical methods***

Mean or medians are given with std. dev. Significance testing of data comparisons was done by standard parametric (Student's t test) and non-parametric (Wilcoxon signed-rank or rank-sum tests) tests or by determining the crossings of confidence boundaries of surrogate datasets (compensated for type I statistical error). Multiple comparisons were corrected using Tukey-Kramer post hoc test. No statistical methods were used to predetermine sample sizes; however, sample sizes were similar or larger than those generally employed in the field. Data collection and analysis were not performed blind to the conditions of the experiments and no randomization was used.

#### MODEL SETUP

The model comprises of two local circuits, one representing the Retrosplenial Cortex (RSC) and the other the Hippocampus (HPC), interconnected through long range projections. The dynamics of neural activity in each of the two regions is described through the mean firing rate of an inhibitory and an excitatory population (Levenstein et al.,2019). The excitatory population of the RSC is subject to a hyperpolarization-activated current $I_h$ (Mehrotra et al.,2023) whereas the excitatory population of the HPC is subject to an adaptive current (Jercog et al 2017; Levenstein et al 2019). This is summarised by the vector equations:

$$\tau_{\mathbf{r}}\dot{\mathbf{r}} = -\mathbf{r} + R_{\infty}(\mathbf{W}\mathbf{r} + \mathbf{b}\mathbf{a} + \mathbf{I} + \zeta_{(\mathbf{t})}) \tag{0.1}$$

$$\tau_{\mathbf{a}}\dot{\mathbf{a}} = -\mathbf{a} + \mathbf{A}_{\infty}(\mathbf{r}) \tag{0.2}$$

where

- $\tau_{\mathbf{r}}$ is a vector whose entries represent the time constant for each population. We set them all equal to 1 for simplicity, time is thus dimensionalised to units of the population rate constant.
- $\mathbf{r} = \begin{bmatrix} r_E^C \\ r_I^C \\ r_E^H \\ r_I^H \end{bmatrix}$ is a vector of firing rates of the excitatory and inhibitory populations of cortex ($r_E^C$,$r_I^C$) and hippocampus ($r_E^H$,$r_I^H$).
- $\mathbf{W} = \begin{bmatrix} W_{EE}^{CC} & W_{EI}^{CC} & W_{EE}^{CH} & 0 \\ W_{IE}^{CC} & W_{II}^{CC} & W_{IE}^{CH} & 0 \\ W_{EE}^{HC} & 0 & W_{EE}^{HH} & W_{EI}^{HH} \\ W_{IE}^{HC} & 0 & W_{IE}^{HH} & W_{II}^{HH} \end{bmatrix}$ is the matrix of connection strengths between populations, such that $W_{EE}^{CH}$ represents the projections strength from population $E$ of region $H$ (Hippocampus) to population $E$ of region $C$ (Cortex). For long-range projections, e.g. originating in HPC and terminating in RSC, we include a 10ms delay transmission to simulate the effects of synaptic transmission between separate brain regions. Note: In this case $W_{EI}^{CH}$,$W_{II}^{CH}$,$W_{EI}^{HC}$,$W_{II}^{HC}$ are set to zero, as we only include long-range projections originating in excitatory neurons in each region.
- $\mathbf{b} = \begin{bmatrix} b_E^C \\ 0 \\ b_E^H \\ 0 \end{bmatrix}$ is the vection of $a$ strengths onto respective populations.

- $\mathbf{a} = \begin{bmatrix} a^C \\ 0 \\ a^H \\ 0 \end{bmatrix}$ is the vector of adaptive current for each population ( $I_h$ for RSC and $a$ for HPC).

- $\mathbf{I} = \begin{bmatrix} I_E^C \\ I_I^C \\ I_E^H \\ I_I^H \end{bmatrix}$ is the vector of background tonic input to each population.

- $\zeta_(\mathbf{t}) = \begin{bmatrix} \zeta_{E,(t)}^C \\ \zeta_{I,(t)}^C \\ \zeta_{E,(t)}^H \\ \zeta_{I,(t)}^H \end{bmatrix}$ is the time-dependent vector of Ornstein-Uhlenbeck (OU) noise which is applied to each population. This is given by:

$$d\zeta = -\frac{\zeta}{\tau}dt + \sigma\sqrt{\frac{2}{\tau}dt}W_t \tag{0.3}$$

  where $W_t$ is a Weiner process. Here we use standard deviation $\sigma = 0.37$ and time scale $\tau = 20$.

- $\tau_{\mathbf{a}}$ is a vector containing the time constants for $a$ in each excitatory population. For simplicity these are identical in both the cortical and the hippocampal population.

- $R_{\gamma,\infty} = g_\gamma[x - \theta_\gamma]_+^2, \gamma = E, I$ represents the threshold-quadratic activation function for each excitatory and inhibitory population. $g$ and $\theta$ are chosen such that $\theta_E < \theta_I$ and $g_E < g_I$, which are necessary conditions for 3 stable states in the phase plane dynamics of each local circuit (Jercog et al.,2017).

- $A_\infty = \frac{1}{1+e^{-k(r-r_0)}}$ represents the sigmoid activation function of $a$ current for each population and depends on two parameters $k$ and $r_0$. Note that for positive $k$ this is an adaptive current which grows during periods of high firing, whereas for negative $k$, this is an inactivity-activated current.

##### ***Model implementation***

Simulations for equations 0.1 and 0.2 are performed in Matlab using the dde23 solver, with OU noise precomputed independently using the forward Euler method with time step dt = 0.01 and a delay of 10ms between inputs arriving via long-range projections from the two regions. Because dde23 uses a variable time step to approximate the solution to the differential equations, we interpolate the solution at every 1ms time-step, making it comparable to sampling frequency of the experimental data. Final simulation statistics are determined for $10^6$

time-steps, in order to have a similar number of UP and DOWN states as the experimental data.

##### *Local Connection Strengths*

The dynamics of the local circuit depend on the particular choices of local parameters $W$ and $I$. To determine the parameters that best fit the dynamics of the biological regions we perform a parameter search across variables $W_{EE}$ (local recurrence) and $I$ (background input). Note: For simplicity, we set $W_{IE} = W_{EE} + 0.2$ and $I_I = I_E - 0.2$. We then compare state duration statistics (UP and DOWN states for CTX and SWR and iSWR states for HPC) for simulations with the experimental data by calculating the Kolmogorov–Smirnov (KS) statistic between the two distributions, as in Levenstein et al., 2019, and define an overall similarity between data and model as:

$$Similarity_{CTX} = (1 - KS_{DOWN})(1 - KS_{UP}) \tag{0.4}$$

and

$$Similarity_{HPC} = (1 - KS_{SWR})(1 - KS_{iSWR}) \tag{0.5}$$

Best parameter fit was defined for parameters that gave the highest similarity between model and data.

##### *Long Range Connection Strengths*

As for local connection strengths, we perform a parameter search in both $E \to E$ and $E \to I$ projection strengths and compare simulation outputs to data. We separately test $RSC \to HPC$ projections and $HPC \to RSC$ projection to verify the causal effect of RSC UP and DOWN states in HPC MUA tonic modulation, and rebound excitation on P(SWR) after a DOWN to UP transition in RSC, as well as the perturbation effect that SWR from the HPC have on the RSC inducing DOWN state transitions. Overall, we select final Long-Range parameters such that they satisfy the following criteria:

- Maintain closest fit to data UP and DOWN state duration statistics.
- Show tonic modulation of HPC MUA by RSC UP and DOWN states, such that HPC MUA is lower during a cortical DOWN state and higher during a cortical UP state.
- Maximise P(SWR) at the UP to DOWN and DOWN to UP transitions.

##### *Model Parameters*

Final model parameters for equations 0.1 and 0.2 are summarised in the following table:

| **Parameter** | **Value** |
|---|---|
| Connection strengths $\begin{bmatrix} W_{EE}^{CC} & W_{EI}^{CC} & W_{EE}^{CH} & W_{EI}^{CH} \\ W_{IE}^{CC} & W_{II}^{CC} & W_{IE}^{CH} & W_{II}^{CH} \\ W_{EE}^{HC} & W_{EI}^{HC} & W_{EE}^{HH} & W_{EI}^{HH} \\ W_{IE}^{HC} & W_{II}^{HC} & W_{IE}^{HH} & W_{II}^{HH} \end{bmatrix}$ | $\begin{bmatrix} 3.04 & -1.5 & 0.13793 & 0 \\ 3.24 & -0.5 & 0.724137 & 0 \\ 0.103448 & 0 & 2.83 & -1.5 \\ 0.068965 & 0 & 3.03 & -0.5 \end{bmatrix}$ |
| $b_E^C, b_E^H$ | $-0.8, 0.8$ |
| $I_E^C, I_I^C, I_E^H, I_I^H$ | 3.35, 2.72, 3.36, 2.77 |
| $\tau_h$ | 100 |
| $g_E, g_I$ | 0.02, 0.05 |
| $\theta_E, \theta_I$ | 0,12 |
| $r_0$ | 2 |
| $k^C, k^H$ | -20, 20 |

**Code availability**

The code used for this study was adapted from the Buzsaki Lab GitHub repository (https://github.com/buzsakilab/buzcode).

**Data availability**

A subset of the data that support the findings of this study will be made available on the Buzsaki lab website, and all data will be made available upon reasonable request.

### Extended Data Figures

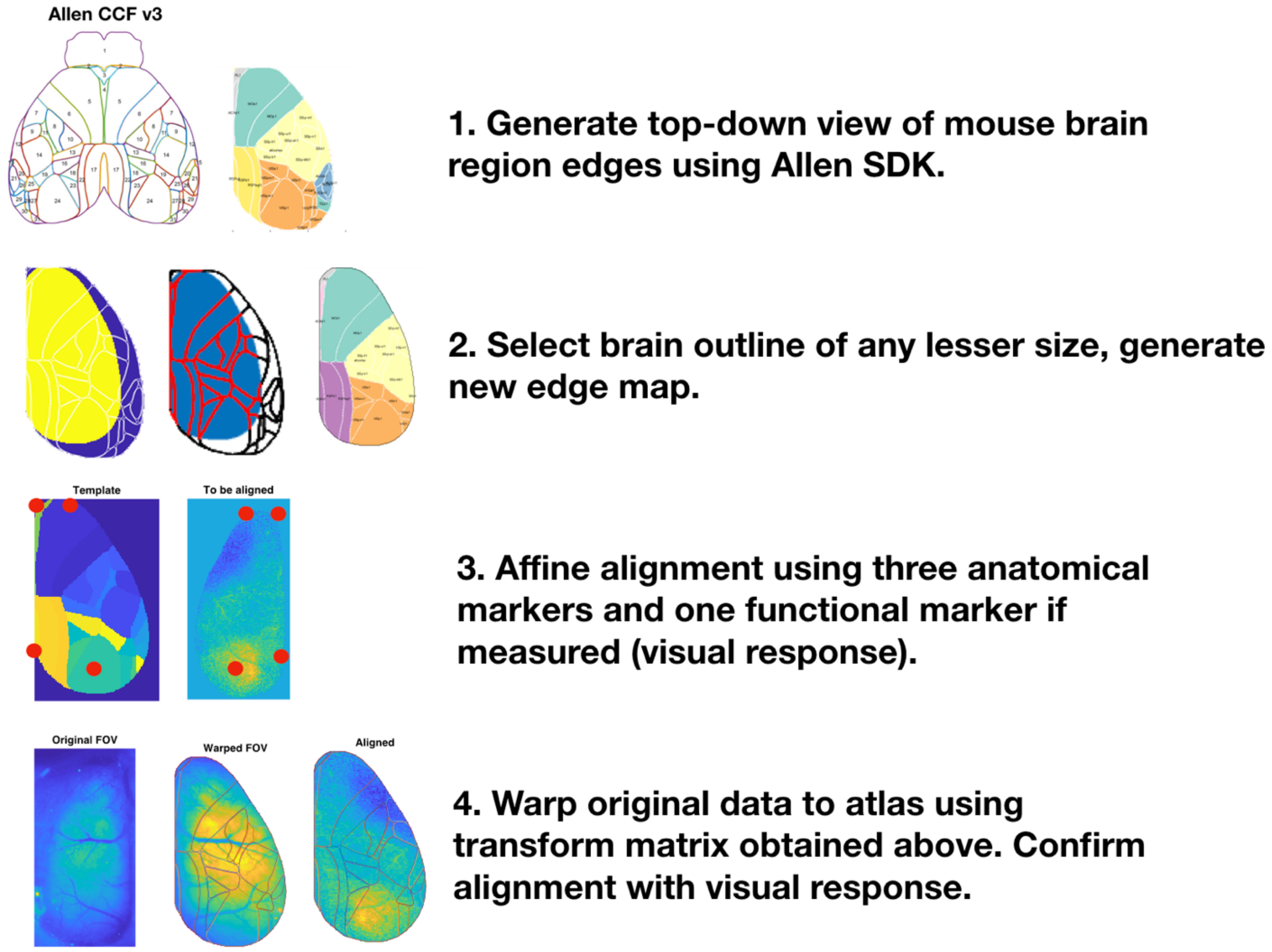


**Supplementary Figure 1,** related to Fig. 1. **Alignment of raw widefield videos to Allen Common Coordinates Framework.** Steps outlined in figure.

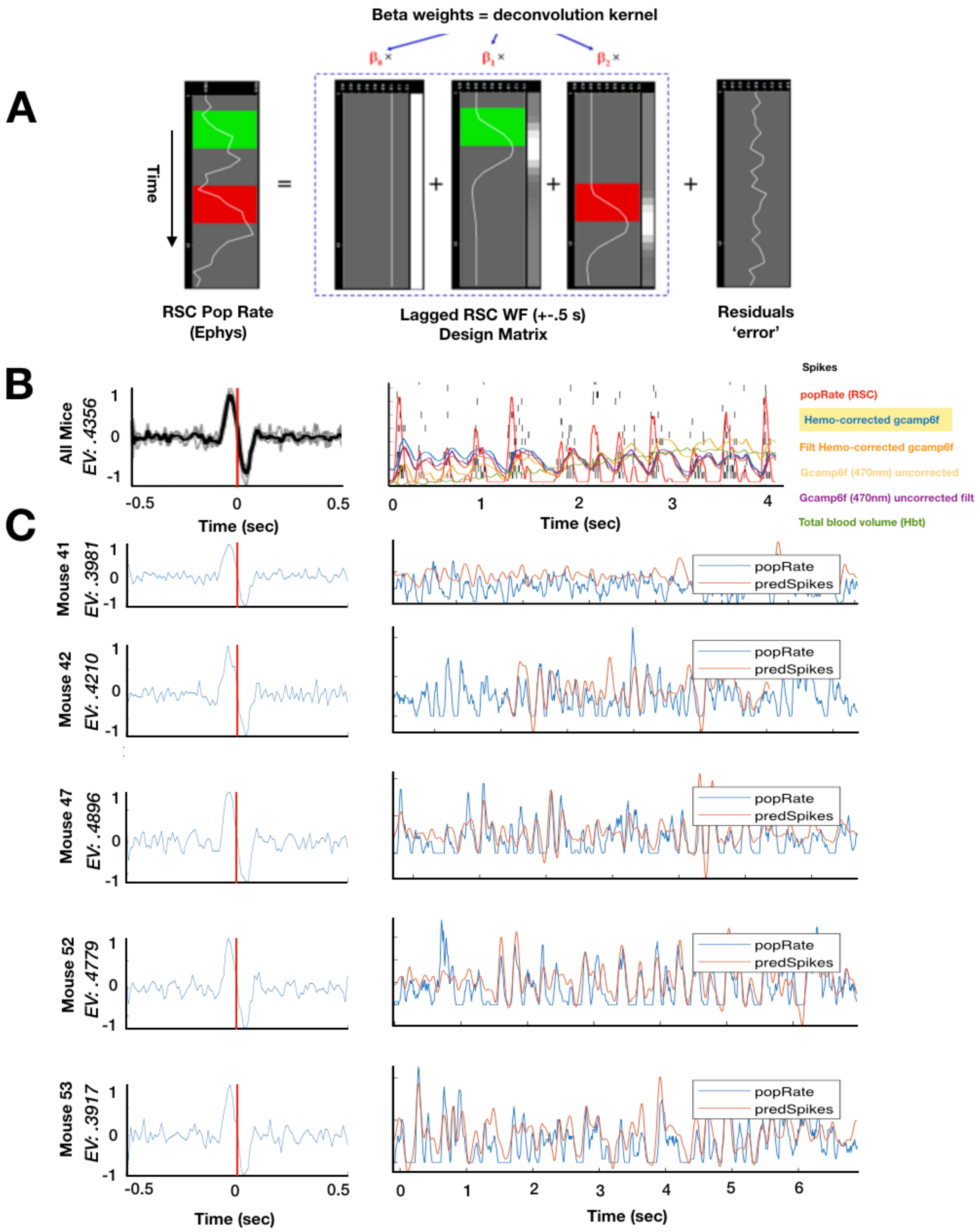

A
Beta weights = deconvolution kernel
Time
RSC Pop Rate (Ephys)
Lagged RSC WF (+-.5 s) Design Matrix
Residuals 'error'
B
All Mice
EV: .4356
Time (sec)
Spikes
popRate (RSC)
Hemo-corrected gcamp6f
Filt Hemo-corrected gcamp6f
Gcamp6f (470nm) uncorrected
Gcamp6f (470nm) uncorrected filt
Total blood volume (Hbt)
C
Mouse 41
EV: .3981
Mouse 42
EV: .4210
Mouse 47
EV: .4896
Mouse 52
EV: .4779
Mouse 53
EV: .3917
popRate
predSpikes
Time (sec)

**Supplementary Figure 2,** related to Fig. 1. **Data-driven deconvolution of widefield data. A.** Schematic of approach. A design matrix is constructed for ridge regression using hemodynamically corrected GCaMP6f data from RSC is lagged ±0.5 seconds, to predict RSC population rate (binned spiking data). The beta weights from each lag are the deconvolution kernel. **B. Left.** Average deconvolution kernel across all mice (black), and individual mice (superimposed gray lines). Total explained variance is 43.6%. **Right.** Representative time series at different stages of preprocessing to illustrate the quality of data and demonstrate the need for deconvolution. Legend on right is colored according to time series it denotes. **C.** Deconvolution results for individual mice and representative time series. Population rate RSC (blue) and predicted rate (orange).

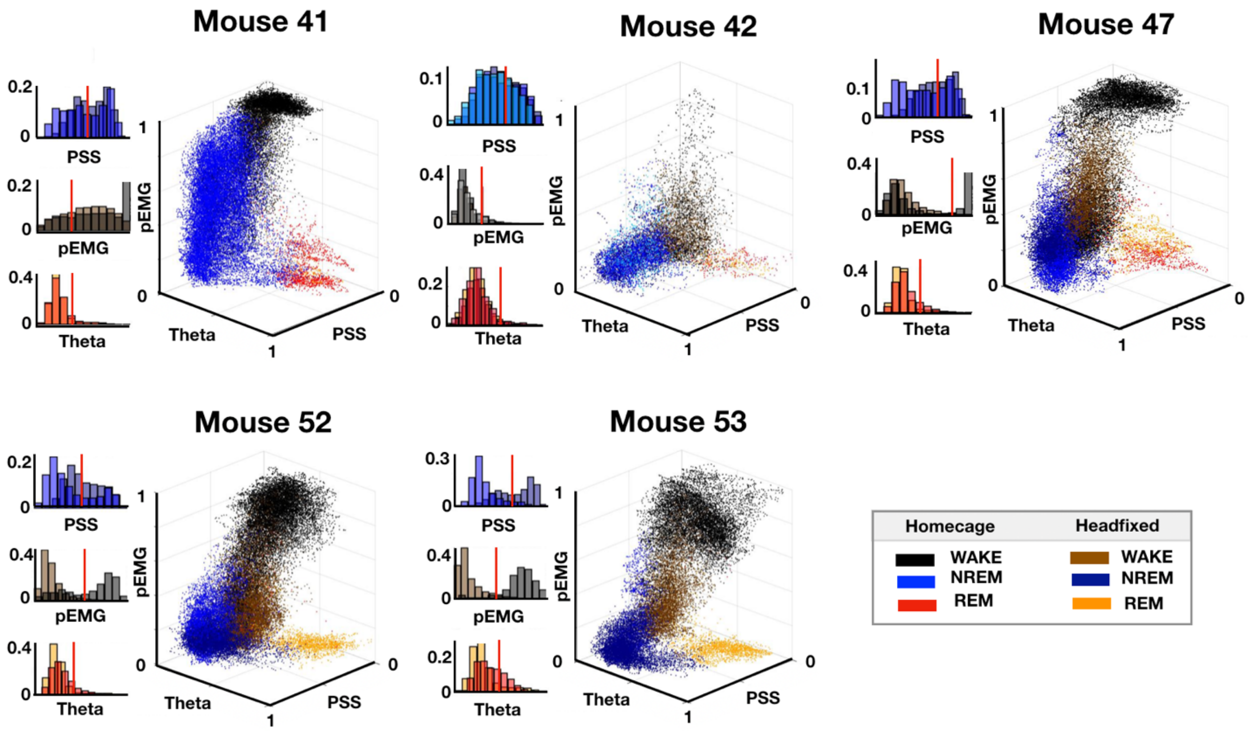


**Supplementary Figure 3**, related to Fig. 2. **Identification of behavioral states**. Comparison of headfixed and homecage sleep in individual mice.

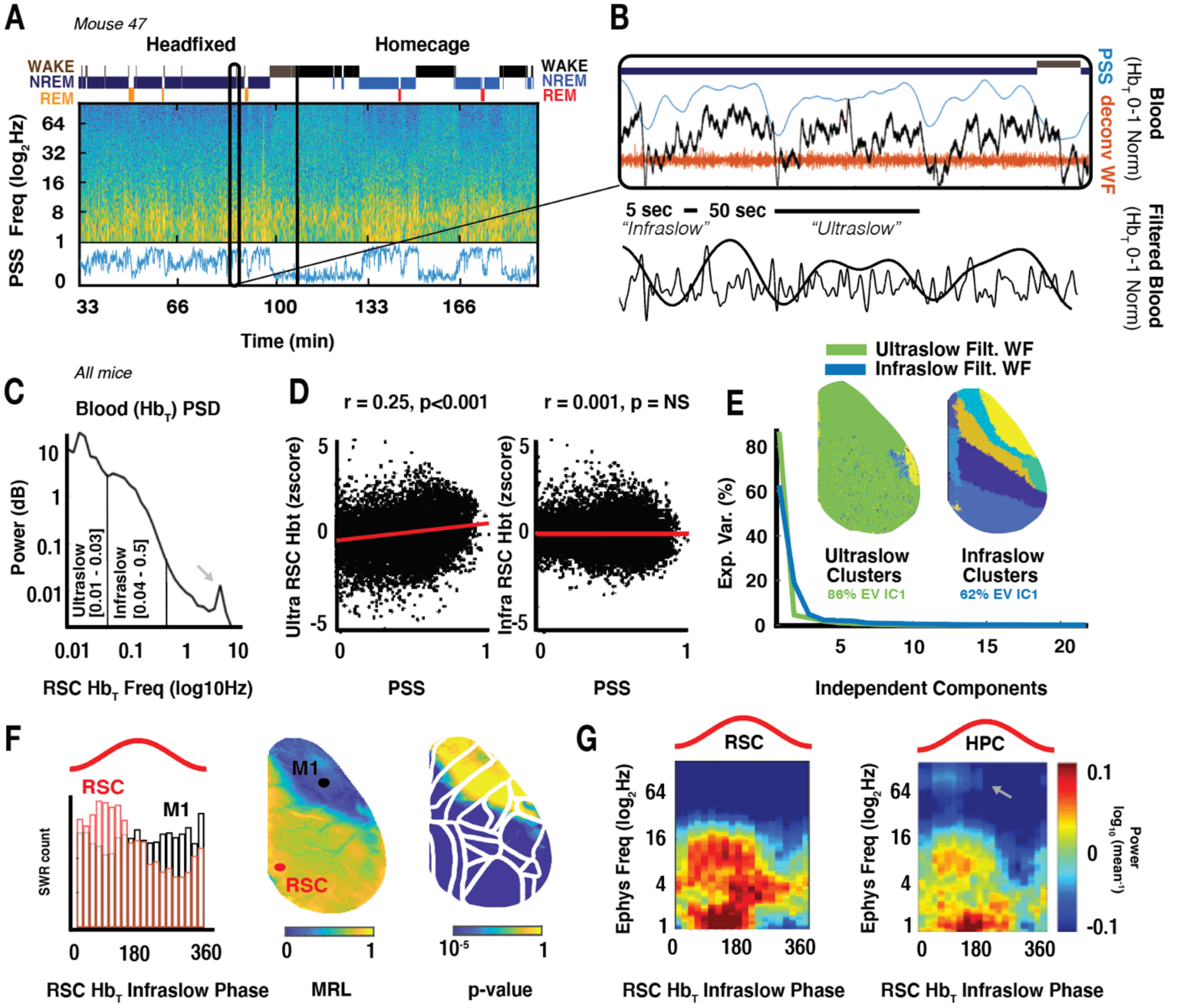


**Supplementary Figure 4,** related to Fig. 2. **Variation in neural state occurs across ultraslow and infraslow timescales. A.** Brain state-scoring of concatenated head-fixed and home cage recording sessions for an example mouse, as in **Fig 2**. Top: WAKE, NREM, and REM brain states, middle: spectrogram of RSC LFP. Bottom: time-varying slope of the power spectrum, PSS, a continuous metric of neural state. **B Top.** Inset in A; plot of PSS (blue), changes in total blood volume (Hbt ch 525; black), and deconvolved widefield activity (orange) within NREM. Note clearly visible ultra- and infraslow changes in total blood volume; **Bottom** Hbt filtered in ultra- and infraslow frequency ranges. Note close correspondence of PSS and ultraslow variation in Hbt. **C.** Average power spectrum of total blood volume (Hbt) across mice. Vertical lines denote ultraslow and infraslow frequency ranges. Arrow, heartbeat. **D.** Spearman correlation between RSC ultraslow-filtered Hbt (**Left**; r = 0.25, $p<0.001$) and infraslow-filtered Hbt (**Right**; r = 0.001*, p = NS*) with PSS. **E.** Independent components analysis (ICA) was performed on both ultraslow and infraslow-filtered Hbt videos. Explained variance for the first 22 independent components is plotted. Note higher explained variance of the first independent component for ultraslow video (86% EV IC1 for ultraslow; 62% EV IC1 for infraslow), corresponding to equal weights across pixels and thus denotes global variation in Hbt. Insets: Average clusters for ultra-

and infraslow-filtered videos, performed using *bz_gradientDescent* (Github repository: Buzcode)*,* a variation of k-means clustering. Initial k = 10. Note smaller number of clusters for ultraslow vs infraslow videos, cumulatively suggesting that ultraslow fluctuations in Hbt are more global than infraslow fluctuations in Hbt. **F. Left.** Distribution of SWRs across infraslow phase in RSC (red) vs M1 (black), leading to high and low MRL values, respectively. **Middle.** Phase modulation (mean resultant length, MRL) of SWRs by total blood volume filtered in infraslow frequency range across pixels. **Right**, Significance map of phase modulation for each pixel (p-values). **G**. Binned spectrograms of RSC (left) and hippocampal (right) LFP during NREM across phases of RSC Hbt infraslow oscillation. Note increased power in delta (1-4 Hz) and spindle (9-15 Hz) bands for both RSC and hippocampus, as well as SWR power in hippocampus (arrow), on the ascending phase and a transition to ~4 Hz oscillation on the descending phase.

**Mouse 41: 31535 DOWNs**

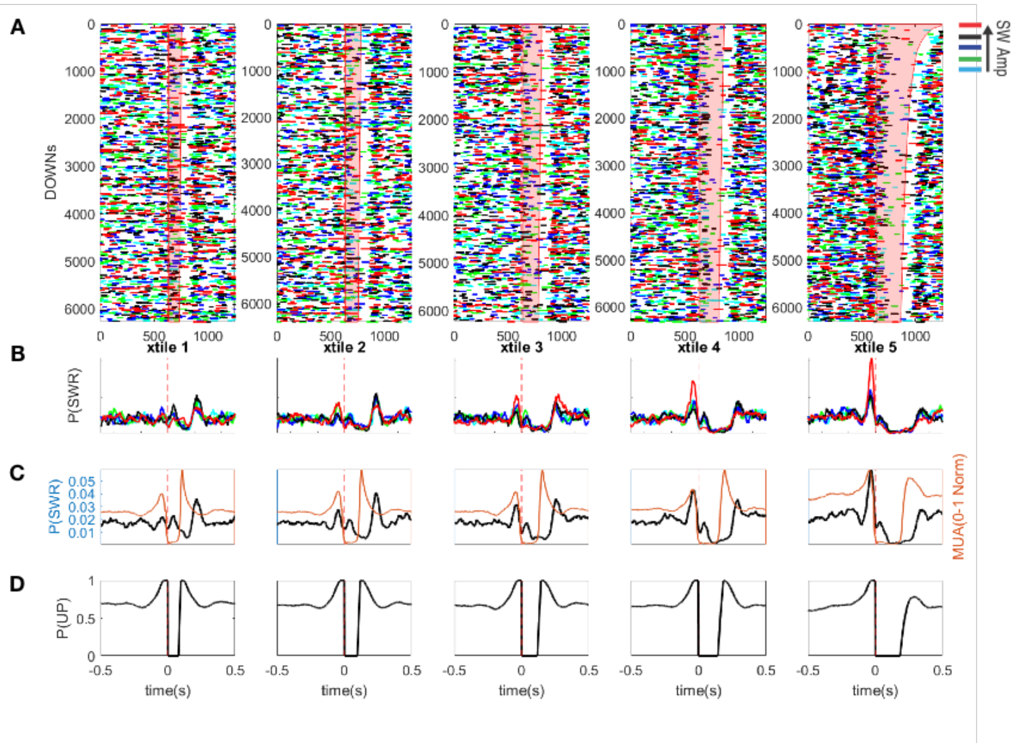


**Mouse 42: 7277 DOWNs**

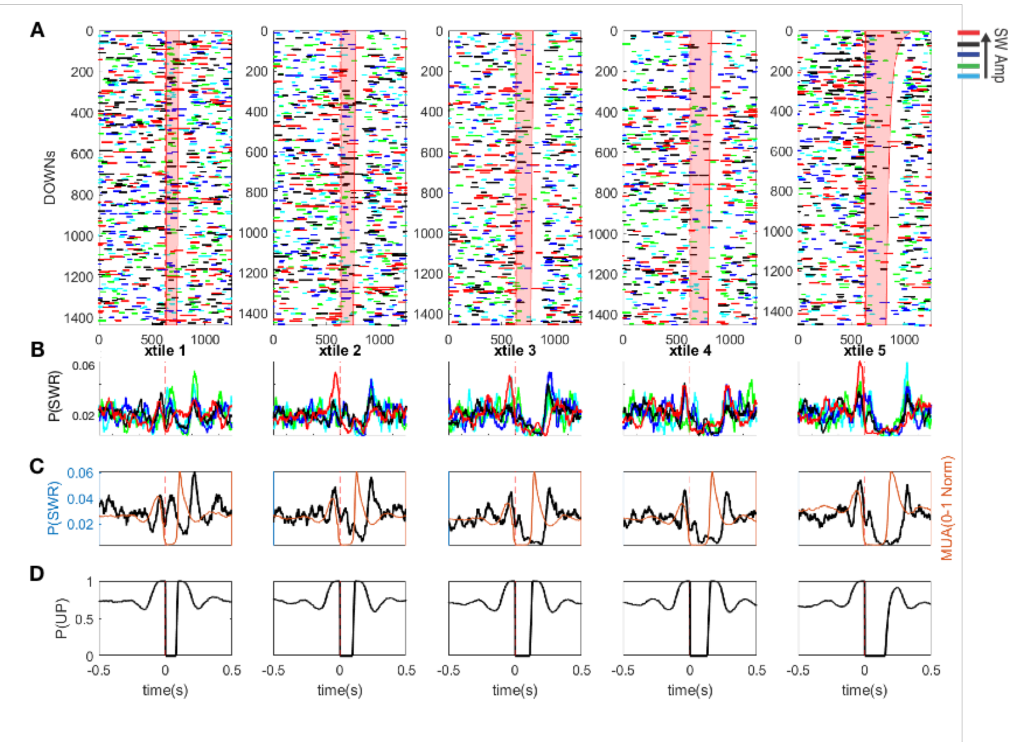

**Mouse 47: 23217 DOWNs**

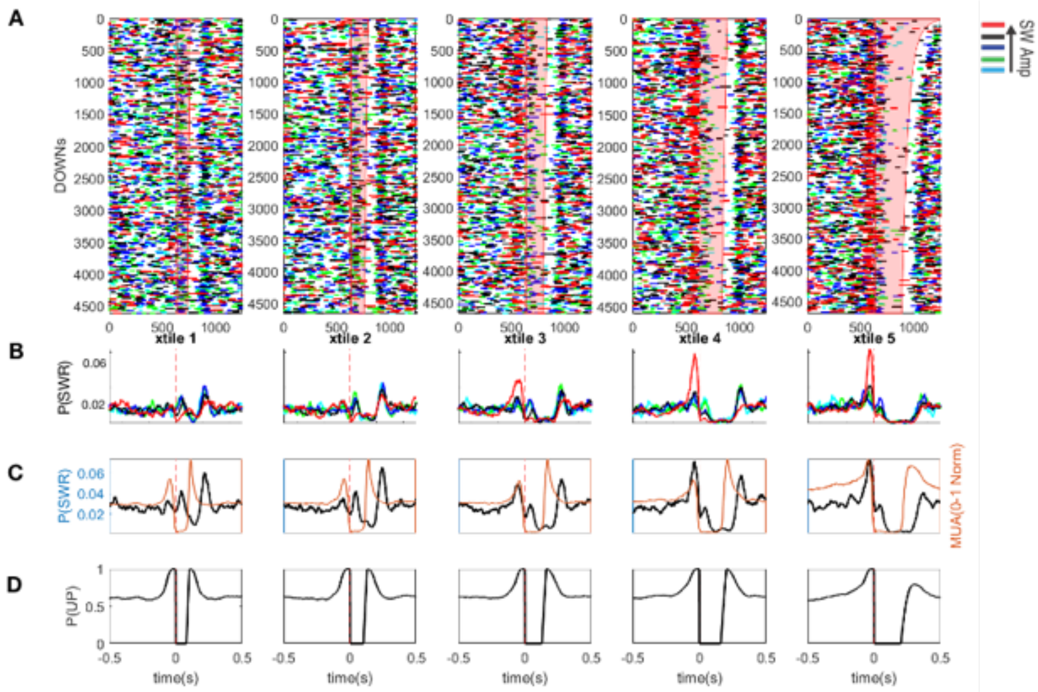


**Mouse 52: 19784 DOWNs**

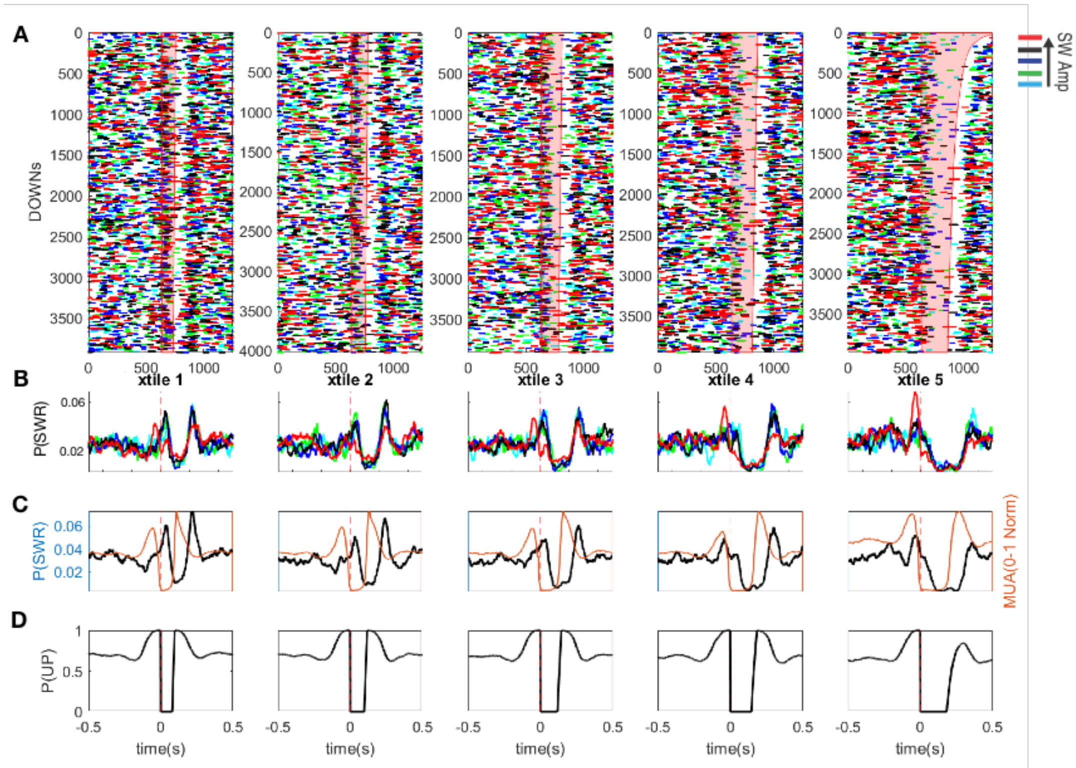

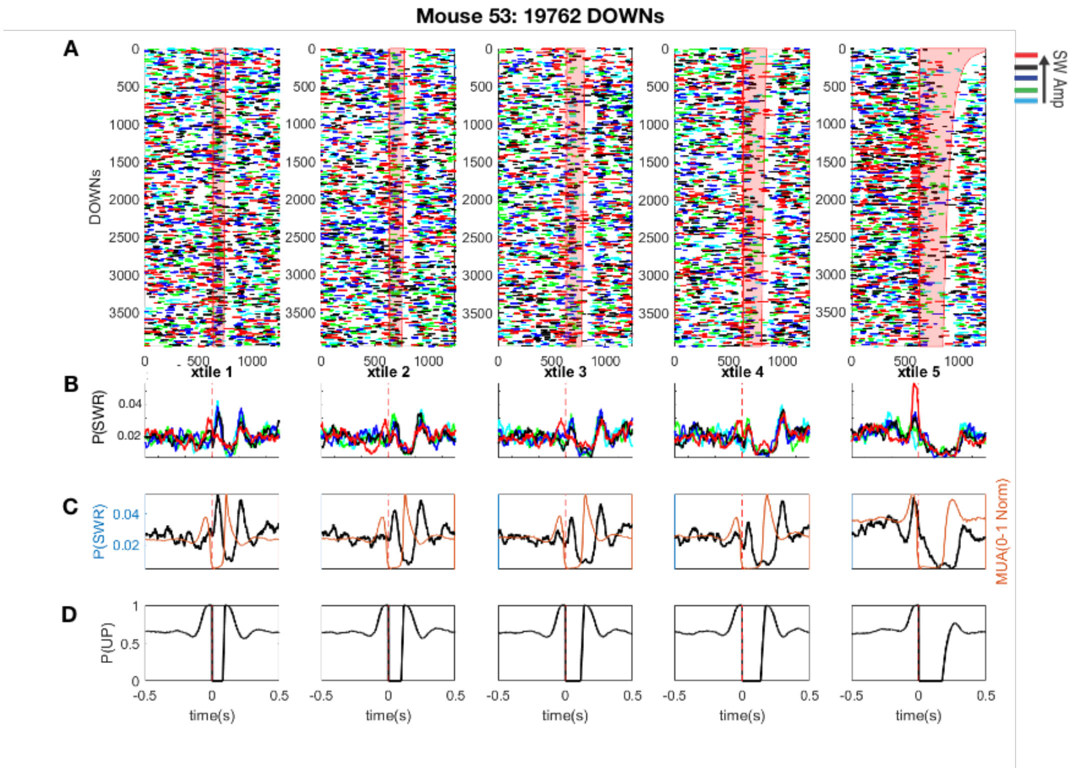


**Supplementary Figure 5,** related to Fig. 4. **Plot of SWR probability surrounding all DOWN states sorted by DOWN state duration quintiles in each mouse. A.** Detected DOWN states split into duration quintiles from short to long (left to right). Raster plot of SWRs surrounding U-D transitions. Line colors denote SWR amplitude; the line length corresponds to the duration of SWR. Pink polygon outlines identified RSC DOWN states (based on RSC MUA). **B.** Probability SWR for each SWR amplitude quintile surrounding U-D transition. Note the increase in large amplitude SWRs with increasing duration DOWN; colors as in A. **C.** Black line P(SWR), orange line (RSC MUA). **D.** Probability of being in an UP state.

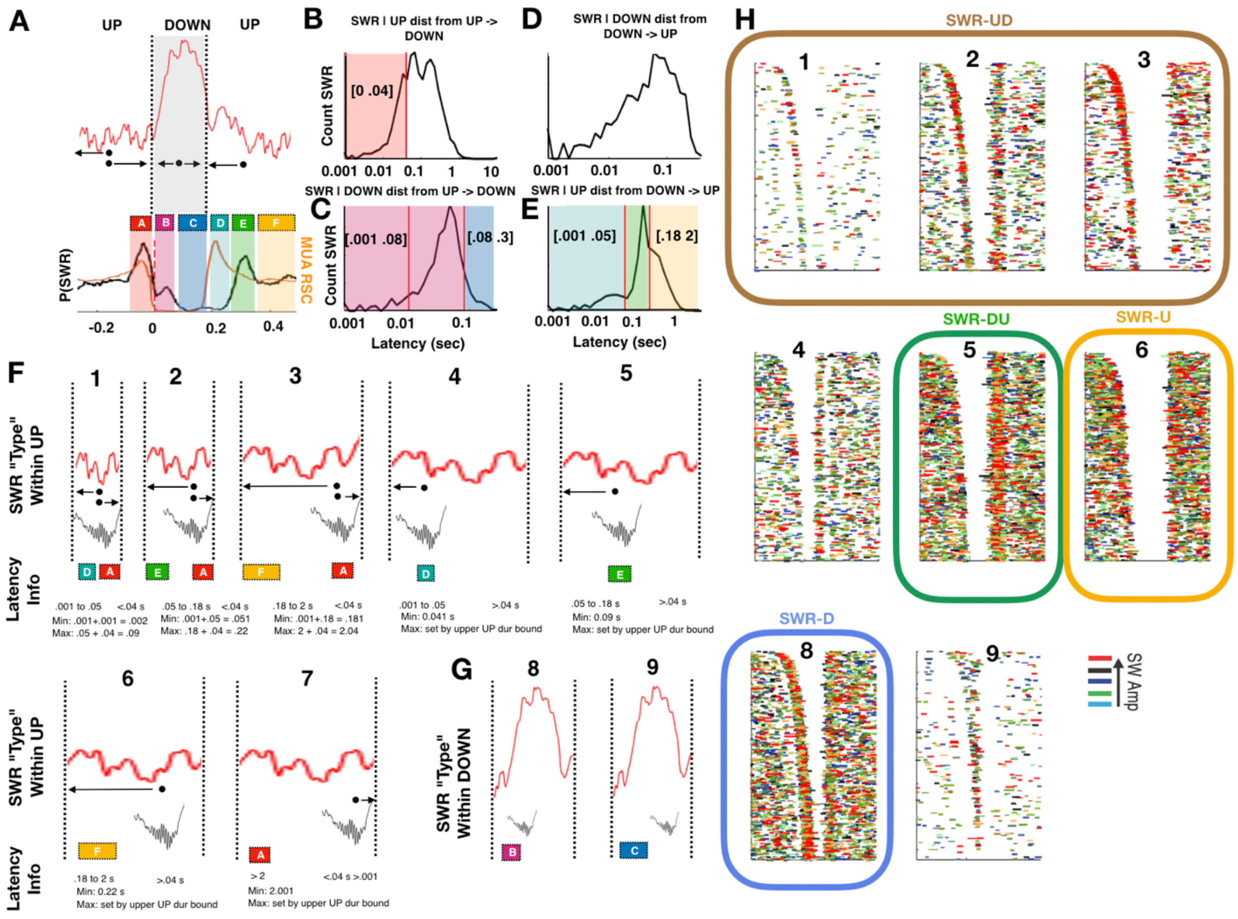


**Supplementary Figure 6,** related to Fig. 4. **Identification of SWR "types" calculated with respect to temporal lag from D-U and U-D transitions in RSC. A.** Schematic of times of interest for determination of SWR types. Top: Red line above denotes UP-DOWN-UP transition. Arrows mark the latencies of interest plotted in B-E (Given in an UP; time to previous D-U, time to upcoming U-D. Given in a DOWN; time to previous U-D, time to upcoming D-U). Bottom: average probability of SWRs (black line) and average RSC MUA (orange line). Letters A-F mark the 'times of interest' used to identify SWR types (colors correspond to plots B-E). **B.** Distribution of latencies of SWRs from U-D transition, given the SWR is in that preceding UP. Red shaded region labeled 'A' defines the interval for $SWR_{UD}$ classification. **C.** Distribution of latencies of SWRs from the most recent U-D transition, given the SWR is in the DOWN state. Purple shaded region labeled 'B' defines $SWR_{D}$ classification. **D.** Distribution of latencies of SWRs to the most recent D-U transition, given the SWR is in the DOWN state. **E.** Distribution of latencies of SWRs to the most recent D-U transition, given the SWR is in the UP state. Blue shaded region labeled 'D' denotes ripples occurring just following the transition. Green shaded region. Labeled 'E' defines $SWR_{DU}$ classification. Yellow shaded region labeled 'F' defines $SWR_{U}$ classification. Note that the distribution of latencies in B-E is not uniform but has clear peaks. These peaks define 'clustering' of SWRs around U-D and D-U transitions, motivating a 'typing' of SWRs. **F,H.** Although SWRs were broken into four types, we highlight a further possible breakdown as a function of the duration of the UP state they are in **F** and **H**. This allows the creation of non-overlapping categories (with the exception of $SWR_{D}$, 50% of which follow the D-U transition @ the same lag as $SWR_{DU}$). However, we ultimately combine SWR 1-3 into one category, $SWR_{UD}$, while noting that there is some overlap between $SWR_{UD}$ and $SWR_{DU}$. **G.** $SWR_D$ definitions and two possible subgroups. **H.** SWR raster plot (lines) surrounding RSC D-U transitions for all DOWN states in an example mouse, colored by SWR amplitude. Numbers are same as in F and G. Outlined boxes correspond to $SWR_{UD}$, $SWR_{DU}$, $SWR_U$, and $SWR_D$ of our final categorization, ultimately used to illustrate the state-dependent bidirectional interaction putatively observed.

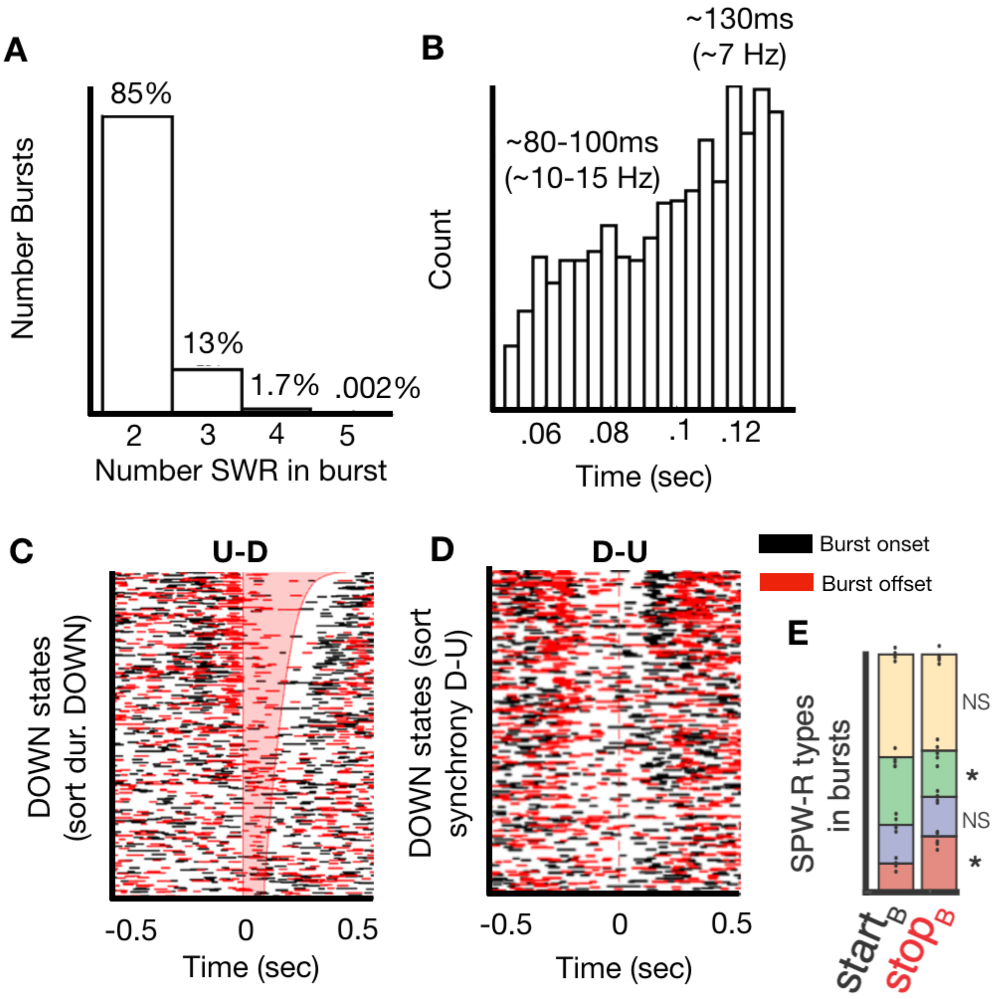


**Supplementary Figure 7,** related to Fig. 4. **Characterization of SWR bursts.** SWR bursts were identified as two or more SWRs with an inter-ripple interval ranging from 50 to 132 ms. **A.** Histogram of identified bursts sorted by number of participating SWRs, note most non-isolated SWRs occur in doublets. **B.** Inter-SWR interval distribution has two distinct peaks, at approximately 7 Hz and 12.5 Hz, likely corresponding to "slow" and "fast" spindles. Both burst frequencies are identified as 'SWR burst' for our purposes. **C-D**. Raster plot of every SWR burst onset (black) and offset (red) for every DOWN state for an example mouse, aligned to U-D transitions (**C**) and D-U transitions (**D**) in RSC. **C** DOWN states, sorted by duration, **D** DOWN states sorted by rebound excitation at the D-U transition. **E.** Proportion of SWR types that can be identified as SWR burst onsets or offsets (color corresponds to SWR type in **Fig 4**). $SWR_{UD}$ (red) were more likely to occur at the end of a burst, SWRDU (green) were more likely to occur at the start of a burst. There was no significant difference between SWRD and SWRU burst onset and burst offset proportion.

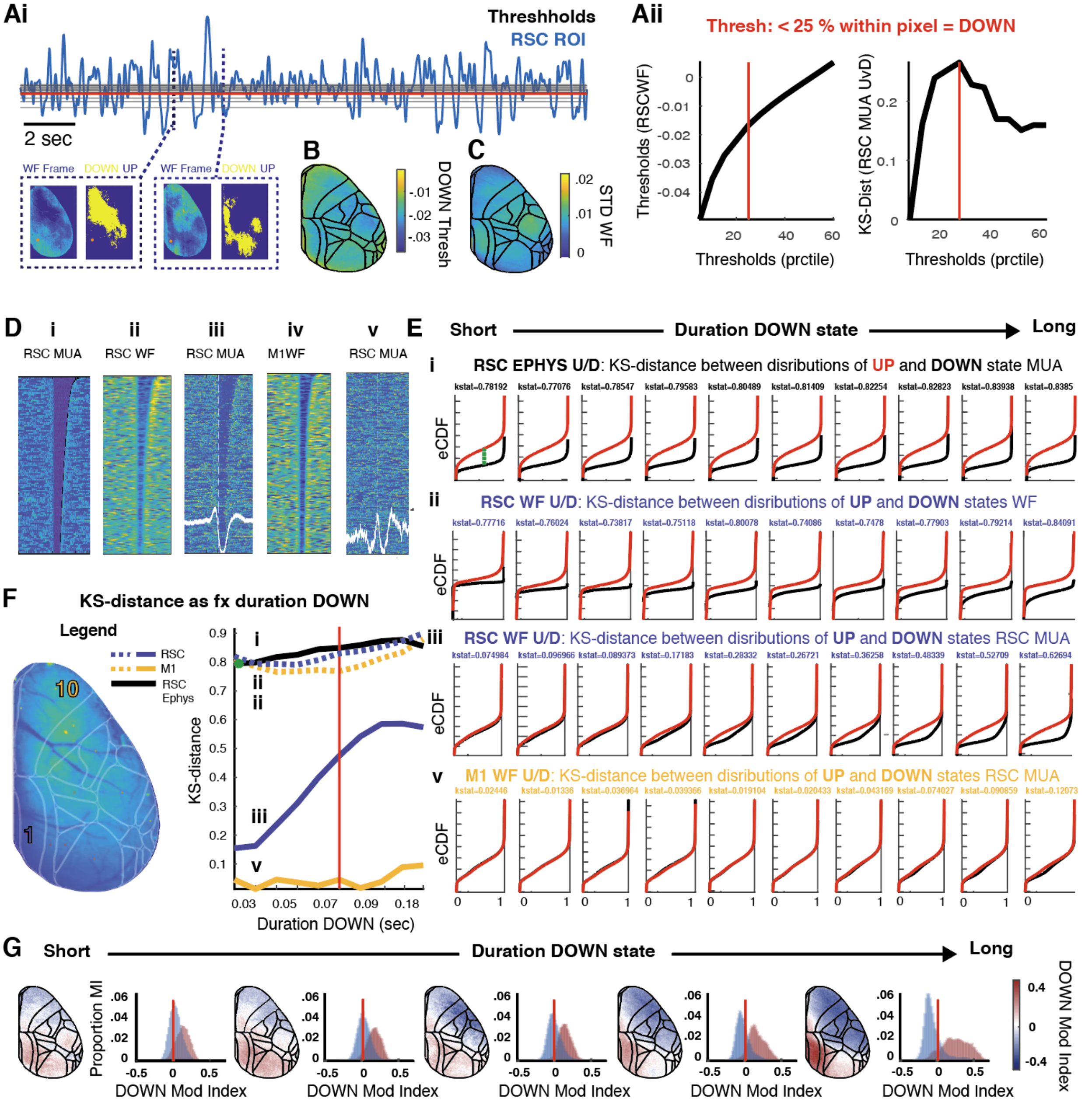


**Supplementary Figure 8,** related to Fig. 6. **Detection of UP and DOWN states in widefield data. Ai-ii.** Identification of the RSC widefield DOWN state threshold that optimally aligns RSC ephys and ophys UP/DOWN states. **Ai.** Deconvolved widefield data from RSC (blue trace) was binarized into putative UP and DOWN states via thresholds that ranged from 5 to 60$^{th}$ percentile (black horizontal lines), above which was classified as an UP state and below as a DOWN state. Vertical dotted lines: example widefield frames plotted at specified time, with the resulting UP/DOWN binarized frame plotted to the right (DOWN states yellow; UP states navy). **Aii, Left.** Widefield values at each percentile threshold. **Right.** Plot of KS distances between distributions of RSC MUA values during WF-detected UP and DOWN states at the specified threshold. Peak separation occurred at the 25th percentile threshold. (red horizontal line, *Ai*, red vertical lines, *Aii*). **B.** Mean 25th percentile threshold detected for each pixel across all videos and mice. Minimal variation across regions. **C.** Standard deviation of detected 25th percentile thresholds across mice for each pixel. **Di.** RSC MUA plotted from -0.5 to 0.5 seconds surrounding U-D transitions detected using ephys data. DOWN states are clearly visible, sorted by duration DOWN state. **Dii.** RSC widefield data plotted -0.5 to 0.5 seconds surrounding U-D transitions detected in that same RSC widefield time series. **Diii.** RSC MUA plotted surrounding those same RSC widefield-detected DOWN states. Qualitative correspondence between Dii and Diii is further quantified in Eii-iii. **Div.** M1 widefield data plotted -0.5 to 0.5 seconds surrounding U-D transitions detected in that M1 widefield time series. **Dv.** RSC MUA plotted surrounding those same M1 widefield-detected DOWN states. Note that there are no visible DOWN states, indicating little or no correspondence between M1 and RSC UP/DOWN states. **Ei-v.** Assessment of DOWN state detection quality as a function of duration DOWN state; using the established 25th percentile threshold. **Ei.** DOWN states are split into duration deciles from short to long, corresponding to plots from left to right, and the empirical cumulative density functions are plotted for MUA values during all UP states within a given duration decile (red), and all DOWN states within the same decile (black). Each plot yields a single value (KS-distance) in plot F. As expected, the value of MUA during RSC DOWN states is smaller than RSC UP states. Also note that regardless of the duration of DOWN state, the distributions remain the same. This indicates the quality of DOWN state detection does not vary as a function of DOWN state duration in ephys data, as expected. To quantify this, the KS-distance between the distributions is computed (green dotted line, leftmost plot), and this distance value is plotted as in **F** (black line, decile 1, mean duration 0.03 sec). The K-S distance is plotted across all deciles, forming a line (E, black line). **Eii-v.** We can now use this to assess the quality of DOWN state detection by asking 2 questions. First, how does DOWN state detection vary by duration DOWN within a widefield region and whether this quality varies between RSC and M1, two selected regions, as shown in **Eii** and **F** (dotted color lines). **Cii.** Same plots as in Ci, but plotting eCDF for distributions of RSC widefield values during RSC widefield-detected UP and DOWN states. This quantifies whether there is a change in quality of detection of DOWN states as DOWN states get shorter for widefield-detected DOWNs. **Ciii.** KS-distance between distributions of UP and DOWN states detected in RSC widefield data, using RSC MUA. This tells us how close the correspondence is between widefield-detected DOWN states and electrophysiologically-defined DOWN states, as a function of the duration of DOWN state. KS-distances for each decile are plotted in **F**, blue line labeled iii. As is expected, detection quality decreases for shorter DOWN states, but above our 0.08 second duration threshold, it is sufficiently consistent for our analyses. **Ev.** KS-distance between distributions of UP and DOWN states detected in M1 widefield data, using RSC MUA. Because there is very little correspondence between M1 and RSC DOWN states, there is little difference between distributions. KS-distances for each decile are plotted in

**F**, gold line labeled v. **F, Left.** RSC and M1 selected for calculation of KS-distances specified. **F, Right.** Plot of KS-distances across duration DOWN state deciles for 2 cases: 1. UP/DOWN KS-distance for DOWN states detected in the time series of interest (dotted lines). All are consistent across DOWN durations and regions, demonstrating similar quality of DOWN detection by region. 2. KS-distance for DOWN states detected in specified region-of-interest (color lines), but computed using RSC MUA. RSC widefield DOWN state-referenced (iii) data against other regions inform us about the spatiotemporal resolution of the widefield data. Taken together, DOWN states can be detected equivalently well across regions, and can be reliably identified for >80 ms events. **G.** Modulation of SWRs by the DOWN states detected in each pixel is plotted in space and as a distribution, from short (left) to long (right) DOWN quintiles. Short duration DOWN states display a unimodal modulation index, but as the DOWN state duration increases, the modulation index becomes bimodal. Blue and red distributions correspond to lateral sensorimotor and default mode networks, respectively, as in Figure 6.

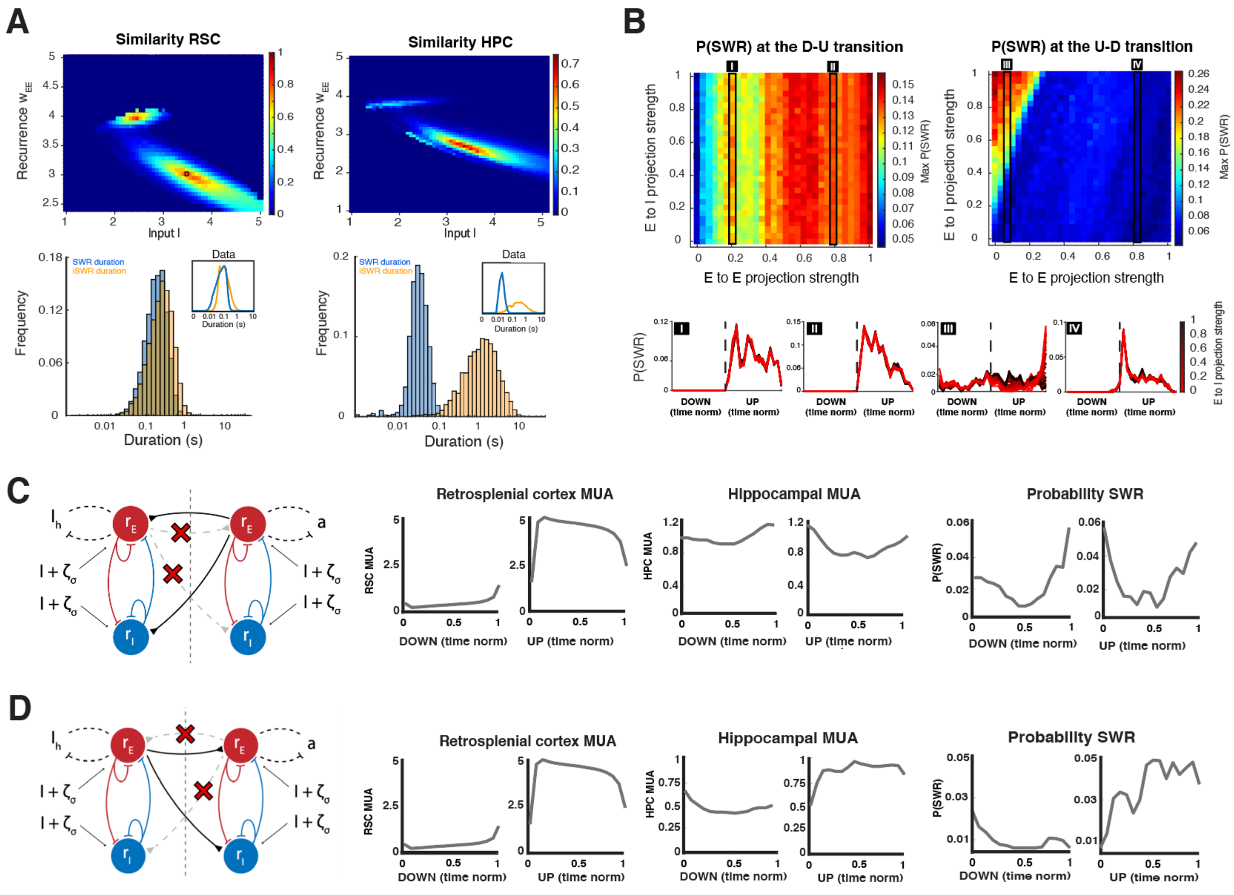


**Supplementary Figure 9,** related to Fig. 8. **Strengths of local and long-range connections are critical for state durations and multi-regional interactions. A. Top.** Local-circuit parameter search space over recurrence in the E populations and tonic background drive; color represents a KS-distance-derived similarity metric (see **Methods**). Bottom. Best-match state duration distributions for simulations and data (insert). **B. Top.** Long-range projection strengths parameter search space over E to E and E to I populations. Left, RSC to HPC projections; right, HPC to RSC projections. **Bottom.** Example P(SWR) for time-normalized RSC UP and DOWN states for each denoted column of the search space (numeric labels I-IV). E-E RSC to HPC projections must be large enough (I) to produce tonic modulation of HPC and a peak in SWR after the D-U transition but not so large that they change the SWR-like dynamics of the HPC (II). HPC to RSC E-E projections must be small enough to show a peak in P(SWR) at the U-D transition (IV vs III), and E-I projections must be large enough (III). **C.** Lesioning the RSC to HPC projections removes tonic modulation of HPC MUA and P(SWR) during UP and DOWN states. **D.** Lesioning the HPC to RSC projections removes the peak in P(SWR) before the U-D transition (compare to figure 8D).

### Supplementary movie legends

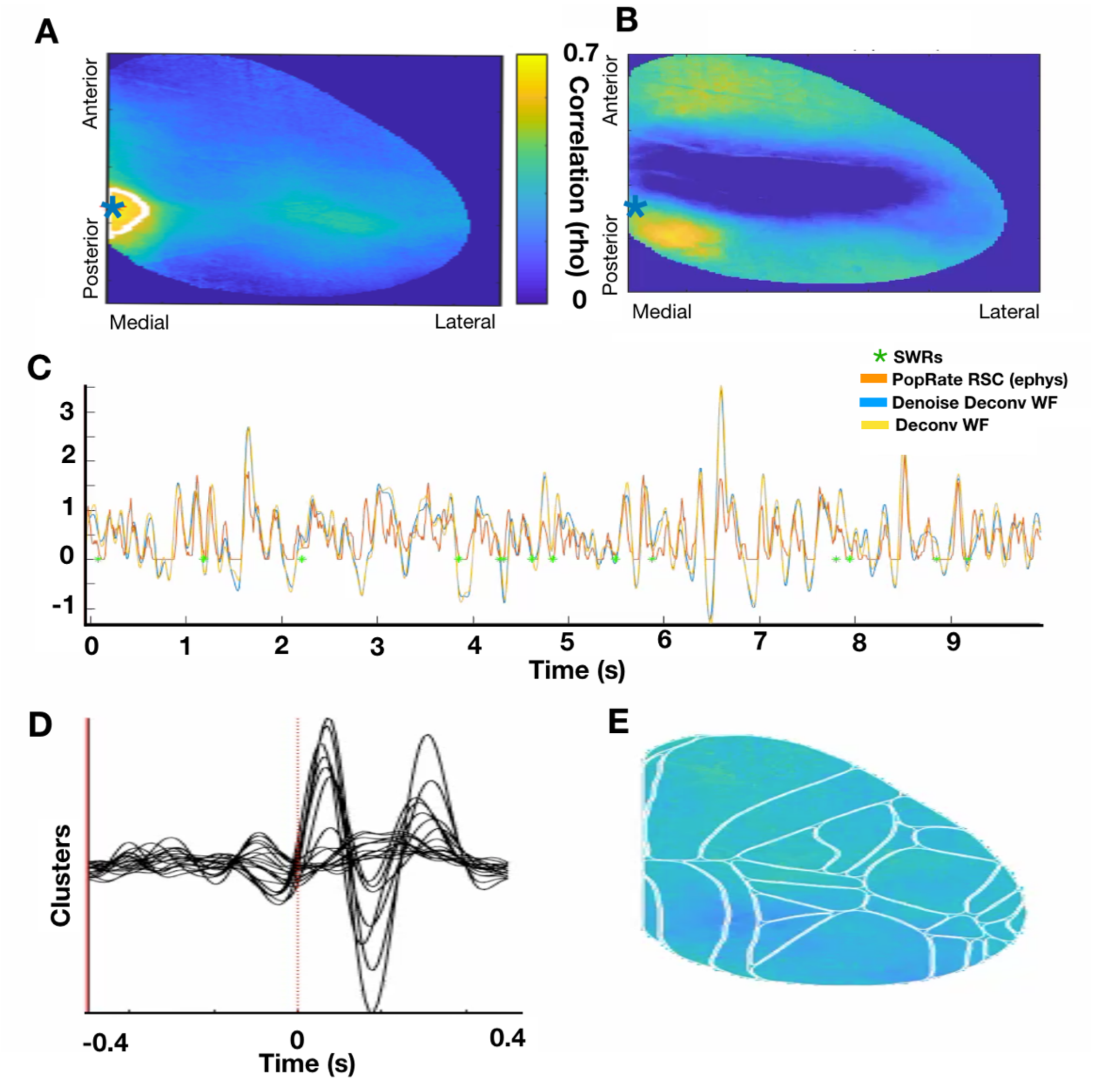


**Supplementary Movie 1. Deconvolution of widefield data** (Still frame shown here). **A.** Pixel-wise correlation between deconvolved widefield time-series and population firing rate demonstrates the highest correlation in RSC (blue star); averaged across all mice. **B.** Video of deconvolved widefield activity corresponding to time series in C. **C.** RSC deconvolved widefield time series from blue star in D (yellow line; blue line: additional smoothing step not ultimately used for data), plotted along with RSC MUA rate (orange). SWRs plotted as green stars. **D.** Average deconvolved widefield response to single presentation of visual stimulus in a V1 pixel across mice. **E.** Movie of single trial responses to visual stimulus in an example mouse. *For movie, click on Supplementary Movie 1ABC* and *Supplementary Movie 1CD.*

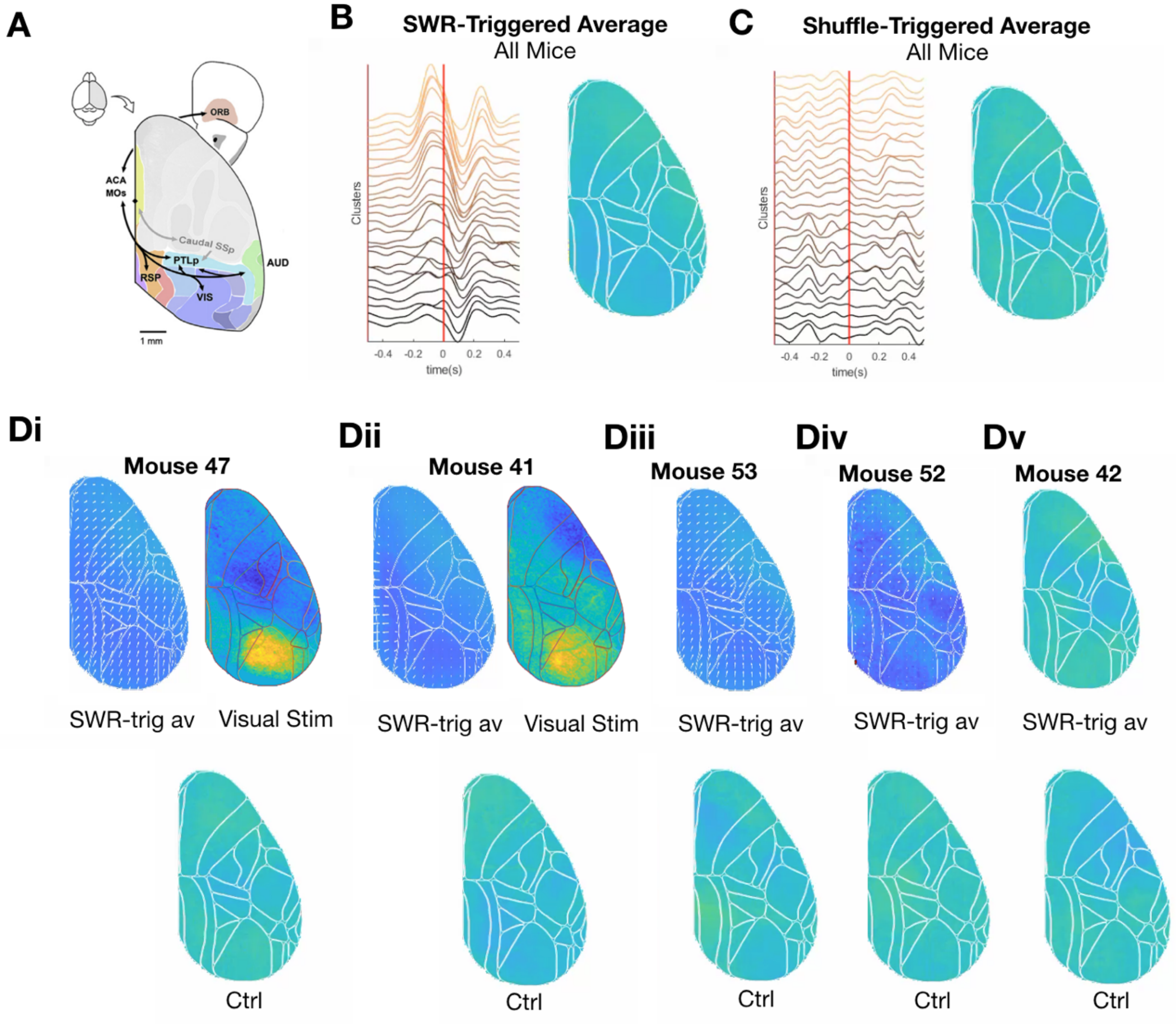


**Supplementary Movie 2. SWR-triggered average movies** (Still frame shown here)**. A.** Anatomical designation of mouse default mode network using cortico-cortical connectivity (Zingg et al., 2014). SWR-triggered average map across all mice. Lines denote average widefield activity from each region, ordered from anterior to posterior (black to orange). **C.** Control: SWR-shuffled average across all mice yields null result, as expected, supporting the notion that these results are not confounded by deconvolution. **D.** Average SWR-triggered widefield activity for all mice (movies), and corresponding visual evoked response in V1 to visual cue for individual mice. **E.** Control triggered-average of shuffled SWR timestamps for each mouse. *For the movies, click on Supplementary Movie 2, and for movies of individual mice, click 47, 41, 53, 52 and 42 .*

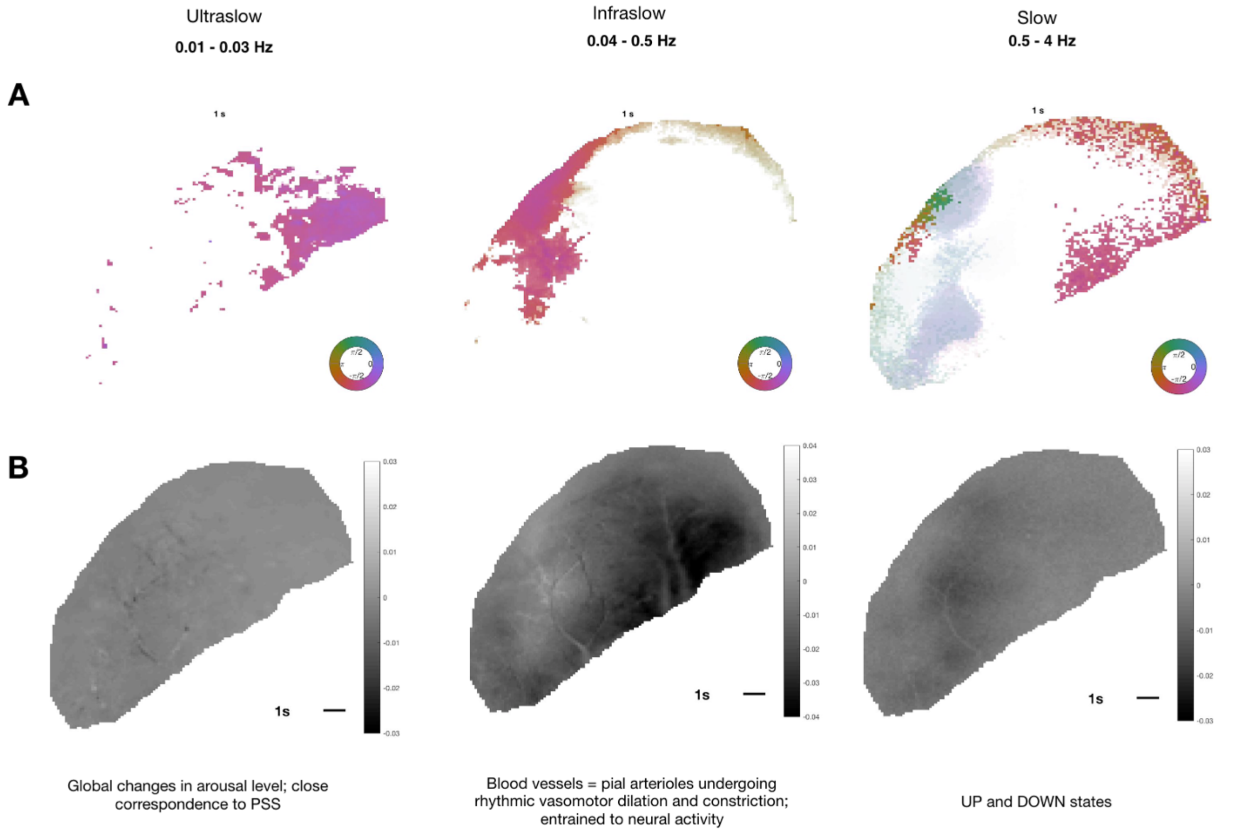


**Supplementary Movie 3. Video of hemodynamically corrected widefield activity filtered in 3 frequency ranges of physiological interest - ultraslow, infraslow, and slow oscillations** *(Still frame shown here)***. A.** Thresholded phase maps of fluorescent signal in a given frequency range, values fade to white with a timelag to display the nature of traveling waves. **B.** Filtered data in the frequency range specified; Ultraslow [0.01- 0.03 Hz], Infraslow [0.04 – 0.5 Hz], Slow [0.5 – 4 Hz]. *For movie, click on Supplementary Movies 3i,ii and iii.*

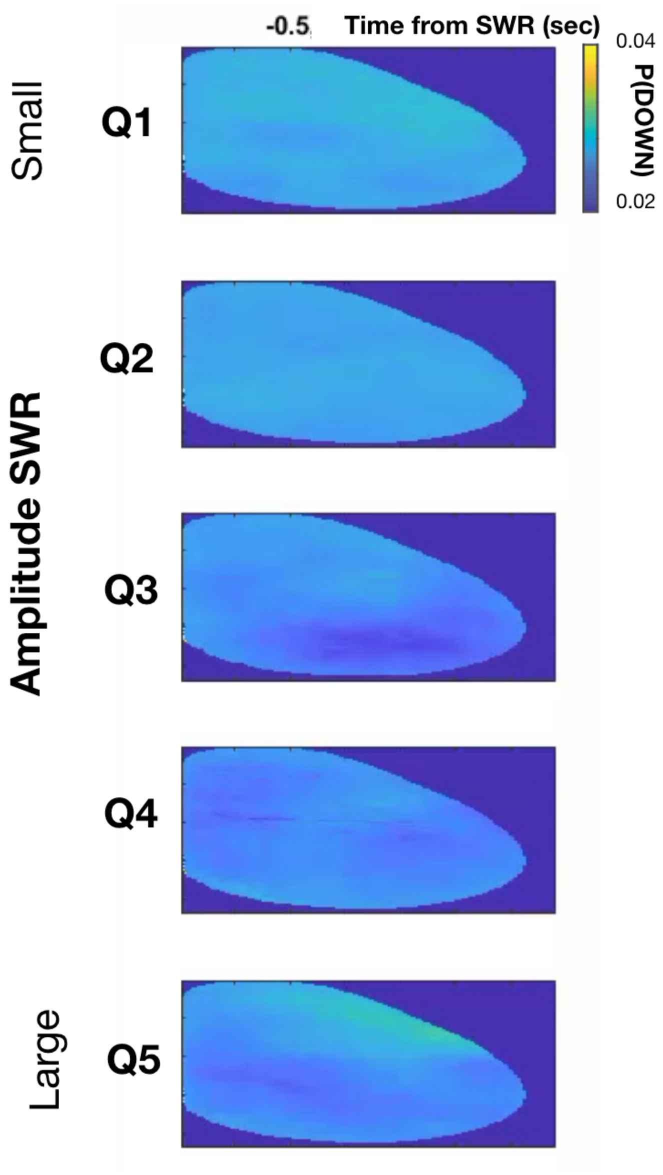


**Supplementary Movie 4. Average probability of DOWN states centered around SWR amplitude quintiles** *(Still frame shown here)***.** Q1-Q5 corresponds to SWR amplitude quintiles, from small to large. T = 0 corresponds to the peak of the SWR. Colormap corresponds to average probability of DOWN state across all mice; all SWRs in the stated amplitude quintile. *For movie, click on Supplementary Movie 4.*

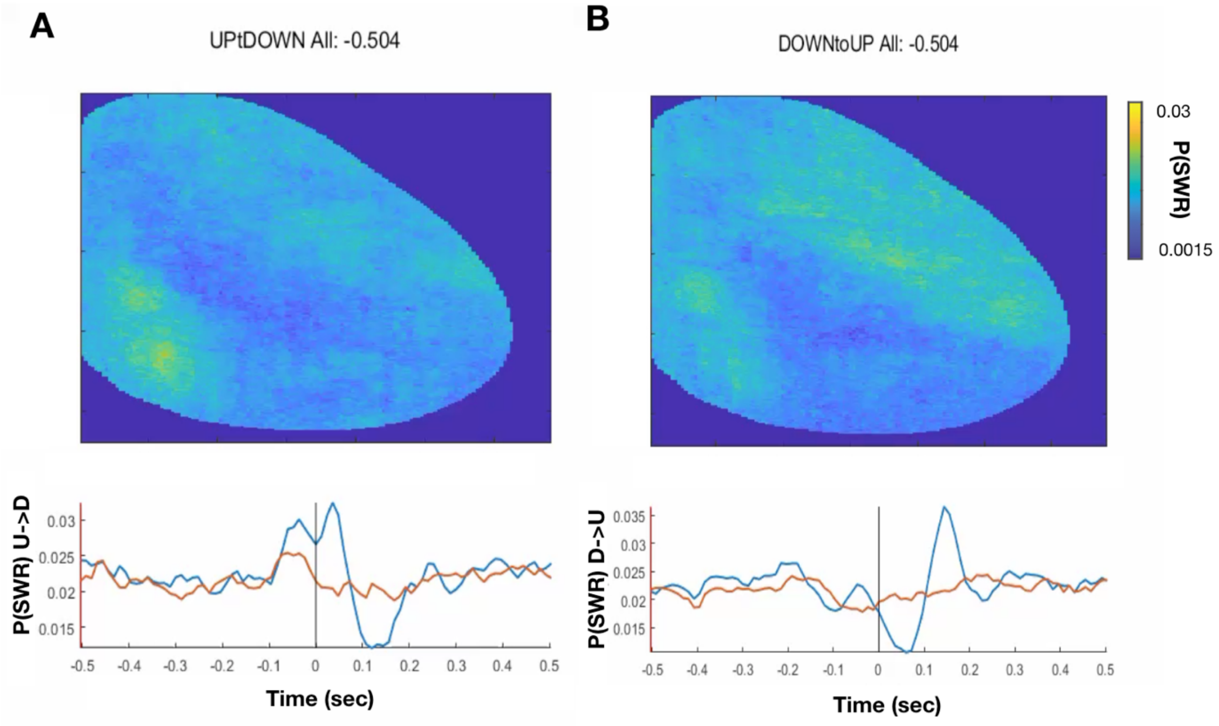


**Supplementary Movie 5. Average probability of SWRs centered at the U-D (A) and D-U (B) transitions, aligned across pixels** *(Still frame shown here)***.** Bottom, Blue line is average P(SWR) surrounding RSC U-D (A) or D-U (B) transition, orange line is average P(SWR) surrounding M1 U-D (A) or D-U. *For movie, click on Supplementary Movie 5.*

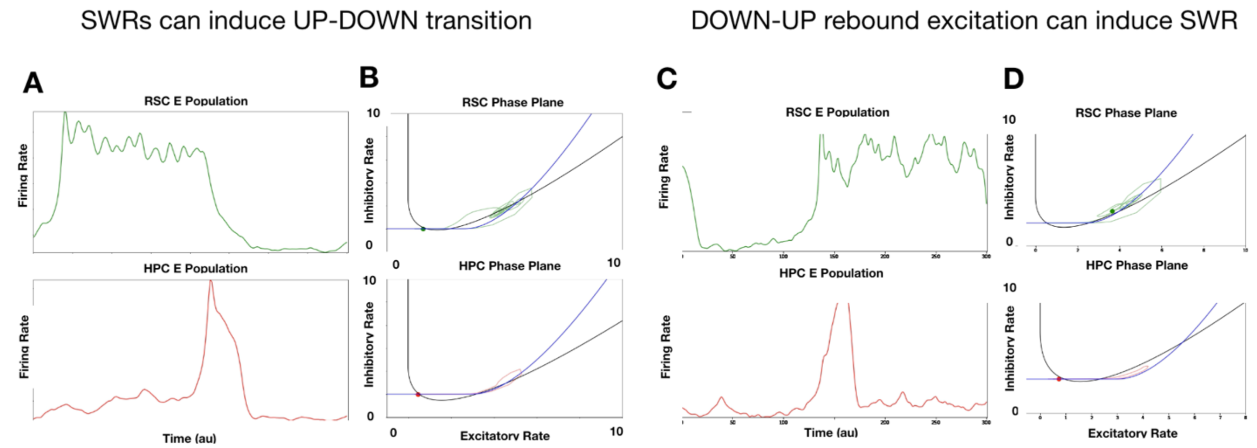


**Supplementary Movie 6. Long-range projections between regions perturb phase plane dynamics causing a state transition. A** and **B**. SWRs in the HPC induce an UP to DOWN transition in the RSC. **A.** Firing rates of the excitatory population in RSC (top) and HPC (bottom) surrounding a SWR. **B.** RSC (top) and HPC (bottom) trajectories on the phase planes corresponding to A. A SWR in the HPC shifts the nullclines of the RSC through long-range projections destabilizing the UP state and causing an UP to DOWN transition. **C** and **D.** Rebound excitation in RSC at the DOWN to UP transition triggers a SWR in the HPC. **C.** Firing rates of the excitatory population in RSC (top) and HPC (bottom) surrounding a SWR, as in A. **D.** As in panel B. H-current build up during a DOWN state produces rebound excitation in RSC (top), displacing E-nullcline in HPC and destabilizing the iSWR steady state and causing a SWR (bottom).
